# The structure of a native orthobunyavirus ribonucleoprotein reveals a key role for viral RNA in maintaining its helical architecture

**DOI:** 10.1101/2021.10.27.466080

**Authors:** Francis R. Hopkins, Beatriz Álvarez-Rodríguez, George R. Heath, Kyriakoulla Panayi, Samantha Hover, Thomas A. Edwards, John N. Barr, Juan Fontana

## Abstract

The *Bunyavirales* order of RNA viruses comprises emerging pathogens for which approved preventative or therapeutic measures for human use are not available. The genome of all *Bunyavirales* consists of negative-sense RNA segments wrapped by the virus-encoded nucleocapsid protein (NP) to form ribonucleoproteins (RNPs). RNPs represent the active template for RNA synthesis and the form in which the genome is packaged into virions, functions that require inherent flexibility. We present a pseudo-atomic model of a native RNP purified from Bunyamwera virus (BUNV), the prototypical *Bunyavirales* member, based on a cryo-electron microscopy (cryo-EM) average at 13 Å resolution with subsequent fitting of the BUNV NP crystal structure by molecular dynamics. We show the BUNV RNP possesses relaxed helical architecture, with successive helical turns separated by ∼18 Å. The model shows that adjacent NP monomers in the RNP chain interact laterally through flexible N- and C-terminal arms, with no helix-stabilizing interactions along the longitudinal axis. Instead, EM analysis of RNase-treated RNPs suggests their chain integrity is dependent on the encapsidated genomic RNA, thus providing the molecular basis for RNP flexibility. Overall, this work will assist in designing anti-viral compounds targeting the RNP and inform studies on bunyaviral RNP assembly, packaging and RNA replication.

**Significance:** Bunyaviruses are emerging RNA viruses that cause significant disease and economic burden and for which vaccines or therapies approved for human use do not exist. The bunyavirus genome does not exist as naked RNA; instead it is wrapped up by the nucleoprotein (NP) to form a ribonucleoprotein (RNP). Using the prototypical bunyavirus, Bunyamwera virus, we determined the 3D structure of the native RNP, revealing a helical architecture with NP molecules linked by lateral contacts only, with no helix-stabilizing longitudinal contacts. Instead, the RNA genome itself plays a role in maintaining the helical architecture, allowing a high degree of flexibility that is critical for several stages of the virus replication cycle, such as segment circularization and genome packaging into virions.

## Introduction

The *Bunyavirales* order of segmented, negative-sense RNA viruses contains over 500 named isolates divided into twelve families (1), five of which include human and animal pathogens. One of these families, *Peribunyaviridae*, comprises arboviruses and is further divided into four genera, with the *Orthobunyavirus* genus being the largest. Important animal-infecting orthobunyaviruses include Schmallenberg virus (SBV) and Akabane virus, which both cause stillbirths and deformities in ruminants, with devastating effects on livestock (2, 3). Important human-infecting orthobunyaviruses include Oropouche, Jamestown Canyon and La Crosse (LACV) viruses, which are associated with acute febrile illnesses with frequent central nervous system infiltration; and Ngari virus, which is responsible for fatal hemorrhagic fever (4–6). Recent reports of the newly-identified Cristoli orthobunyavirus and the re-emergence of Umbre orthobunyavirus as agents of fatal encephalitis, confirm the risk posed by orthobunyaviruses to human health (7, 8). Whilst vaccines have been developed against several animal-infecting orthobunyaviruses, including SBV (2, 9), no FDA-approved vaccines or therapies are available for those infecting humans. As insect habitats shift in response to climate change, arboviruses pose a growing threat and present a pressing need for new strategies in vaccine and anti-viral design.

Bunyamwera virus (BUNV) is the prototypic orthobunyavirus that serves as a tractable model for the *Bunyavirales* order as a whole. BUNV virions are pleiomorphic and include a host-derived lipid envelope coated in virus-encoded glycoprotein spikes (10) that surrounds the tri-segmented RNA genome comprising small (S), medium (M) and large (L) segments. Each RNA segment is encapsidated by multiple copies of the nucleocapsid protein (NP) to form a ribonucleoprotein (RNP), which associates with the virus-encoded RNA-dependent RNA polymerase (RdRp; L). RNPs serve as the active templates for all viral RNA synthesis activities and represent the form in which the RNA segments are packaged into virions.

The BUNV NP comprises a globular core with N- and C-terminal lobes bisected by a positively-charged RNA-binding groove (11–16) that accommodates approximately 11 RNA bases. The N- and C-termini protrude from the core to form ‘arms’ that mediate homotypic oligomerisation, and this overall arrangement is shared by NPs from SBV, LACV and Leanyer orthobunyaviruses (11–16). Orthobunyavirus NP crystallises as closed, non-physiological, tetrameric or hexameric rings (16) and this oligomeric promiscuity is confirmed by mass spectrometry and electron microscopy (EM), which reveal oligomers of increasing size from trimer to octamer (14, 17, 18).

Previous EM analyses of native RNPs released from virions show they are highly flexible and have resulted in apparently conflicting models that describe their overall architecture (Figure S1). As its name suggests, the ‘beads on a string’ model proposes RNPs contain NP monomers interacting with immediately adjacent neighbours through lateral contacts only, which is based on EM observations of flexible RNPs with a diameter of single NP monomers (13, 14). The alternative model proposes that RNPs adopt a relaxed helical architecture, which is supported by EM observations of RNPs that exhibit a staggered positioning of monomers with a cross-section of two NPs, which was interpreted as a helix of ∼4 NPs per turn (11, 12); and also by the crystallization of RNA-free LACV NP oligomers in a helical array (12).

Native RNPs from the *Arenaviridae*, *Phenuiviridae* and *Nairoviridae* families within the *Bunyavirales* order also possess a flexible architecture (19–21). In contrast, a high-resolution cryo-electron microscopy (cryo-EM) model of a rod-like RNP from a member of the *Hantaviridae* family reconstituted from recombinant protein, revealed a compact and rigid helical architecture, maintained by both lateral and longitudinal NP-NP interactions through the helix (22). However, to date, no high-resolution structural information from native RNPs exists for any member of the *Bunyavirales* order.

We present a model of native BUNV RNPs extracted from infectious virions, based on a cryo-EM average at 13 Å resolution and subsequent fitting of crystal structures by molecular dynamics. The gross structural features of the RNP were determined and our model suggests that the helical arrangement is not maintained by longitudinal NP-NP interactions, but is dictated by lateral NP-NP interactions and the encapsidated RNA. This work will aid the design of anti-viral compounds specific to the orthobunyaviral RNP and inform future studies into the mechanisms of viral processes such as RNP assembly and packaging.

## Materials and Methods

### RNP Purification

BUNV was propagated in BHK-21 cells and purified as described previously (23, 24) and RNPs were purified based on a previous protocol (11) (SI Materials and Methods). Centrifugal concentrators were used for buffer-exchange of pure RNPs to facilitate EM analysis.

### Negative staining EM

Purified virions or RNPs were adsorbed to glow-discharged carbon coated copper grids, stained with 1% uranyl acetate and imaged in a 120 kV FEI Tecnai 12 microscope.

### Cryo-EM and Image Processing

Cryo-EM and cryo-electron tomography (cryo-ET) data were collected on an FEI Titan Krios EM 300 kV microscope (Astbury Biostructure Laboratory, University of Leeds). Collection and on-the-fly data pre-processing was performed as previously described (25). Cryo-ET reconstruction and sub-tomogram averaging (STA) were performed respectively using IMOD (26) and PEET (27). Cryo-EM data was processed by a single particle approach using Relion (28) to produce a final symmetrical reconstruction at 13 Å (SI Materials and Methods).

### Molecular dynamics

A ‘split-NP’ (an NP monomer containing the N- and C-terminal arms from its neighbours) was fitted into the EM average using helical symmetry and molecular dynamics were used to refine the fit using MDFF (29) (SI Materials and Methods).

### Mini-genome replication assay

BUNV RNPs were reconstituted as previously described (30, 31). Briefly, BSR-T7 cells in 24-well plates were transfected with 0.2 μg each of pT7riboBUNL, pT7riboBUNN or mutant derivatives, pT7riboBUN-SREN and 0.1 μg pCMV-Firefly-Luc as a transfection control. At 24 h post-transfection, luciferase expression was measured using the dual-luciferase assay kit (Promega).

### RNase treatment

Purified BUNV RNPs were diluted in TNE buffer (100 mM Tris-HCl, 200 mM NaCl, 0.1 mM EDTA, pH 7.4, supplemented with protease inhibitors (Roche)) and mixed with indicated amounts of RNase A to a final volume of 20 μL. Samples were incubated for 30 minutes at room temperature and visualised by negative staining EM.

## Results

### BUNV RNP purification

To structurally characterise native BUNV RNPs, virion purification and RNP isolation was optimised. BUNV was purified from infected cell supernatants by ultracentrifugation, with purity assessed by SDS-PAGE (Figure S2A) and negative staining EM (Figure S2B), which revealed intact as well as disrupted virions from which viral RNPs were released. RNPs were purified from freeze-thawed virions by ultracentrifugation (adapted from (11)) after testing different methods to release RNPs (Figure S3), with RNP-containing fractions detected by western blotting (Figure S4A). Peak fractions corresponded to approximately 23% OptiPrep and subsequent analysis by negative staining EM revealed abundant, filamentous RNPs (Figures 1A and S4B). The width of all RNPs was uniform along their length and corresponded to two NP monomers. These monomers adopted a staggered arrangement consistent with a helical conformation (Figure 1A, inset), and there was no indication of a ‘beads on a string’ morphology that would result from single monomers joined only by lateral contacts. Any disruption to the helical conformation was only apparent at the sharpest (≥90°) bends in the filament. Of note, almost all observed RNPs were circularized and linear filaments were rare (less than one percent of the imaged RNPs). Although the RNA itself might not be resolved by negative staining, we expected to visualise a region along the RNP where the NP organisation was different, distinguishing the segment ends from non-terminal regions. No such region common to all segments was identified. However, some RNPs contained an associated density potentially compatible with that of the viral RdRp (Figure S5). These densities lack the additional globular domains appended to the ring-like structures reported for RdRps from negative-sense RNA viruses (32, 33) but they do resemble recent images of reconstituted polymerase-associated RNPs from an arenavirus (34). Different buffers were explored to optimise RNP separation. No discernible difference in the overall helical RNP appearance was observed, although we noted that TNE with 25 mM NaCl resulted in the straightest RNPs that would be most amenable to EM processing (Figure S6).

**Figure 1.**
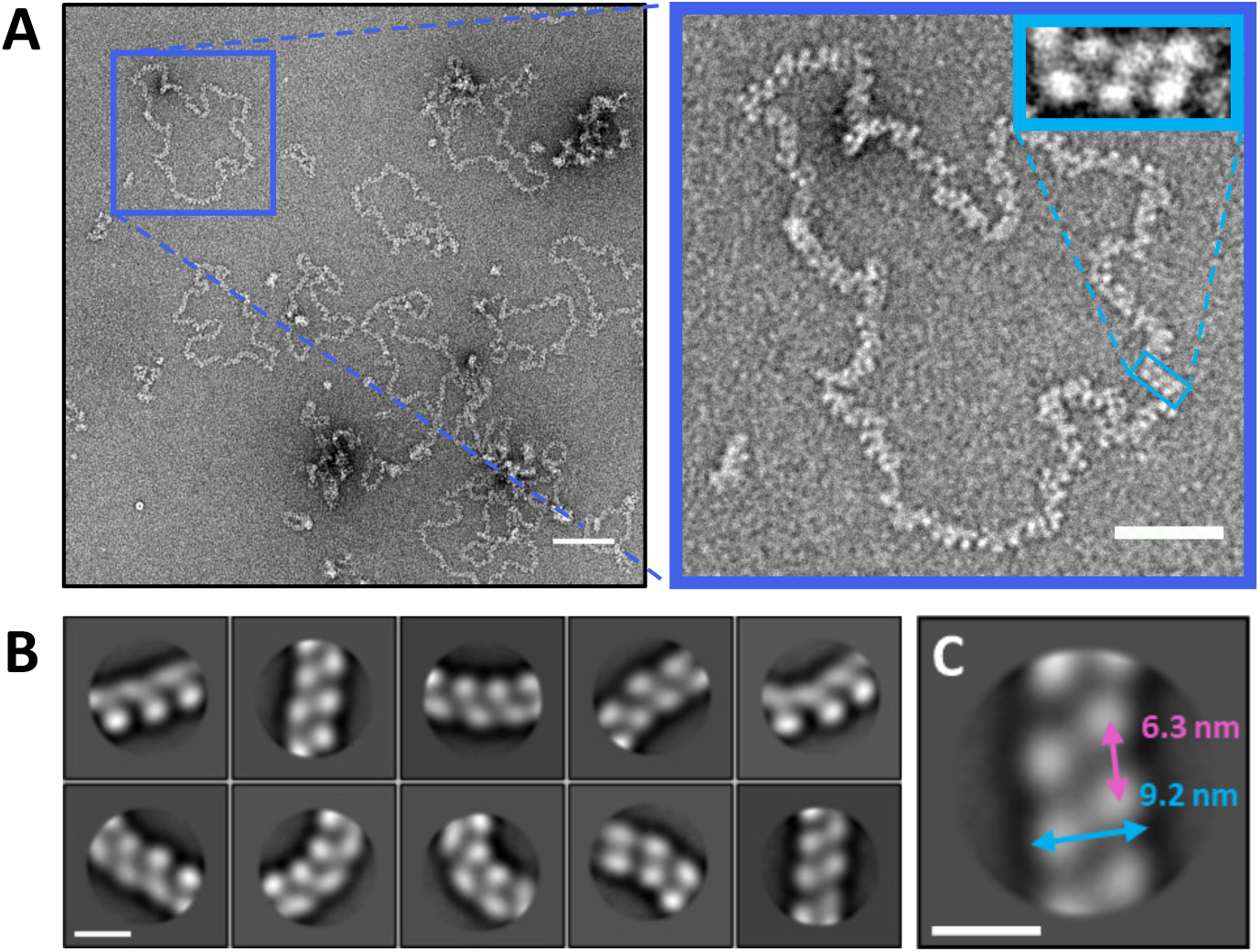
Negative staining EM analysis of purified BUNV RNPs. **(A)** Negative staining EM confirmed the abundance and purity of RNP filaments, and regions of apparent helical organisation were apparent (inset). Scale bar: 100 nm in main and 50 nm in inset. **(B)** Alignment and classification clearly showed helix-like architecture within filaments. Scale bar: 10 nm. **(C)** Preliminary estimation of the RNP width and helical pitch. Scale bar: 10 nm.

### BUNV RNPs have a helical NP organisation

To gain further insight into the organisation of the BUNV RNPs, we measured their lengths. Given the nucleotide lengths of BUNV RNA segments (961 nt for S; 4458 nt for M, and 6875 nt for L), and that the orthobunyaviral NP is approximately 5 nm wide and binds approximately 11 RNA bases (11–15, 35), the ‘beads on a string’ organization would predict their lengths to be around 430, 2020 and 3125 nm, respectively. Approximately 600 discrete, circularized RNPs were manually traced and measured (Figure S7A), with the majority falling into three distinct species with mean lengths of approximately 160, 650 and 1050 nm, suggesting a consistent degree of condensation (Figure S7B). Assuming a helical NP arrangement, the helical rise was estimated to be ∼1.7 nm, by dividing the RNP lengths by the estimated number of NP monomers per segment based on the NP:nucleotide stoichiometry.

Alignment and 2D classification of particles picked along the RNP was used to further elucidate the architecture of the filament. As RNP filaments exhibited a high degree of flexibility, which allowed them to bend up to ∼90° angles, particles were picked with a relatively small box size of 294 Å (Figure S4B). Alignment and 2D classification of these particles clearly showed a pattern of alternating NP monomers along the filament length, consistent with a helical structure (Figure 1B). Analysis of straight class averages allowed measurement of the width of the filament and of the distance between monomers along the length of the filament, which in principle represents an approximation of the helical pitch. Measuring three points (centre and each end) along the filament in the ten class averages (Figure 1B) resulted in mean values of 9.2 nm for the width and 6.3 nm for the longitudinal distance between monomers (Figure 1C). This corresponds to ∼3.7 monomers per helical turn (6.3 nm helical pitch over 1.7 nm helical rise, as determined above). We note that the value for the helical pitch varied from 4.9 to 8.6 nm, in agreement with the observation that the distances between individual monomers can vary to accommodate bends in the filament.

### Cryo-electron microscopy produces a 13 Å average of a native BUNV RNP

To generate a 3D map of the BUNV RNP, we adopted an approach inspired by that of Bharat et al., (36, 37) which combined cryo-ET and cryo-EM to study helical filaments. The use of a Volta phase plate during data collection resulted in high-contrast tomograms in which clear RNP filaments were apparent (Figure 2A). Additionally, large electron dense particles not present in the negative staining preparations were observed that possibly represented condensed RNPs within the air-water interface, but did not obstruct the reconstruction of tomograms or the sub-tomogram selection along the RNP lengths. STA of RNPs was performed using a cylindrical reference and a 20 Å average was produced, which clearly displayed the helical nature of the filament, with a width of ∼76 Å and helical pitch of ∼65 Å (Figures 2B and S8). As EM data does not have a definitive handedness, we imaged the RNPs using AFM (Figure S9), which can be used to determine the handedness of helical assemblies (38). Right-handed RNP sections were imaged in standard and high-resolution AFM (Figures S9A inset, and S9B). The handedness of the RNPs was more apparent when averaging RNP sections using a right-handed reference (Figure S9C). Of note, the average was also right-handed when using a reference with an inverted hand (Figure S9D). Therefore, we concluded BUNV RNP helices are right-handed.

**Figure 2.**
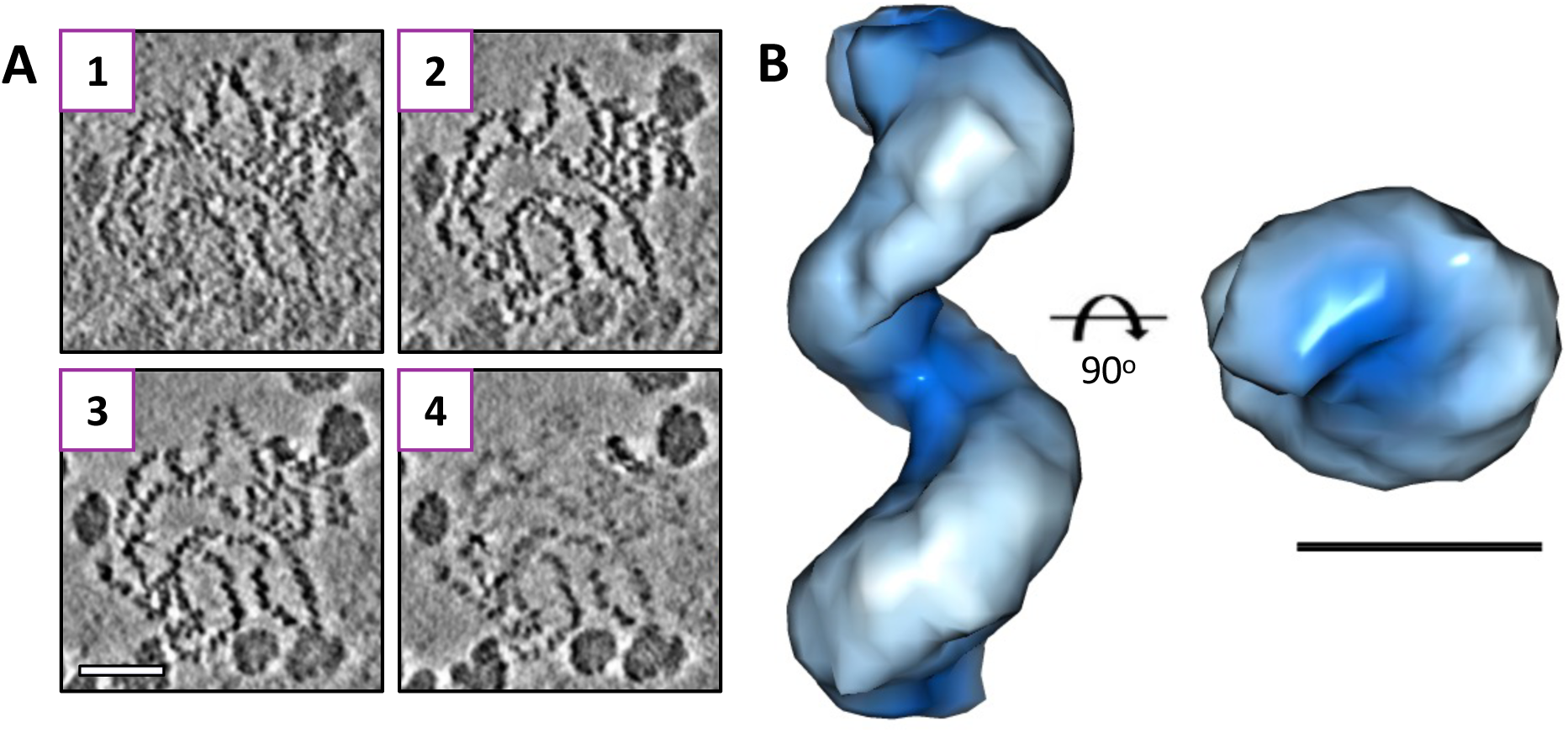
Cryo-ET derived model of a BUNV RNP. (**A**) Reconstructed tilt-series revealed clear RNPs and numerous electron dense particles. Insets 1 to 4 show slices 13.5 Å apart from within an area containing RNP(s). Scale bar: 50 nm. (**B**) Sub-tomogram averaging produced a 20 Å reconstruction of a helical BUNV RNP. Scale bar: 5 nm.

A cryo-EM data set was then collected with Volta phase resulting in ∼135,000 straight RNP particles (Figure S10). Given the absence of long straight RNP segments, particles were selected and processed using a single particle approach. To validate the helical nature of the RNP, these particles were first subjected to 3D classification using a cylinder as a reference (removing any possible bias towards a helical model). The resulting averages were clearly helical (Figure S11), and therefore the particles were taken forwards for image processing using the STA-derived model. Following a similar approach as described recently for influenza virus (39), which involved classifying particles into small sets of ∼4,000 homogeneous particles, we obtained a 14 Å resolution average without symmetry applied (Figures S12A and B). When applying helical symmetry to this average, with searches centred on -100° for twist and 15 Å for rise, the searches routinely converged on -104.9° twist and 17.7 Å rise. Therefore, these numbers were applied for a final reconstruction with local searches, resulting in a 13 Å average that converged on -105.26° twist and 18.25 Å rise (Figures 3A and S12C). The final symmetrised map is very similar to the non-symmetrised map (cross-correlation of 0.94) and to the map generated using a cylindrical reference (cross-correlation of 0.87), suggesting the helical parameters we have applied are correct. In this average, the helix appears not to be tightly wound and compact, with the distances between neighbouring rungs of the helix appearing too large for stabilising contacts, which would facilitate the flexibility observed within the filaments. It is also clear in this model that the number of monomers per turn (3.7) is not a whole integer.

**Figure 3.**
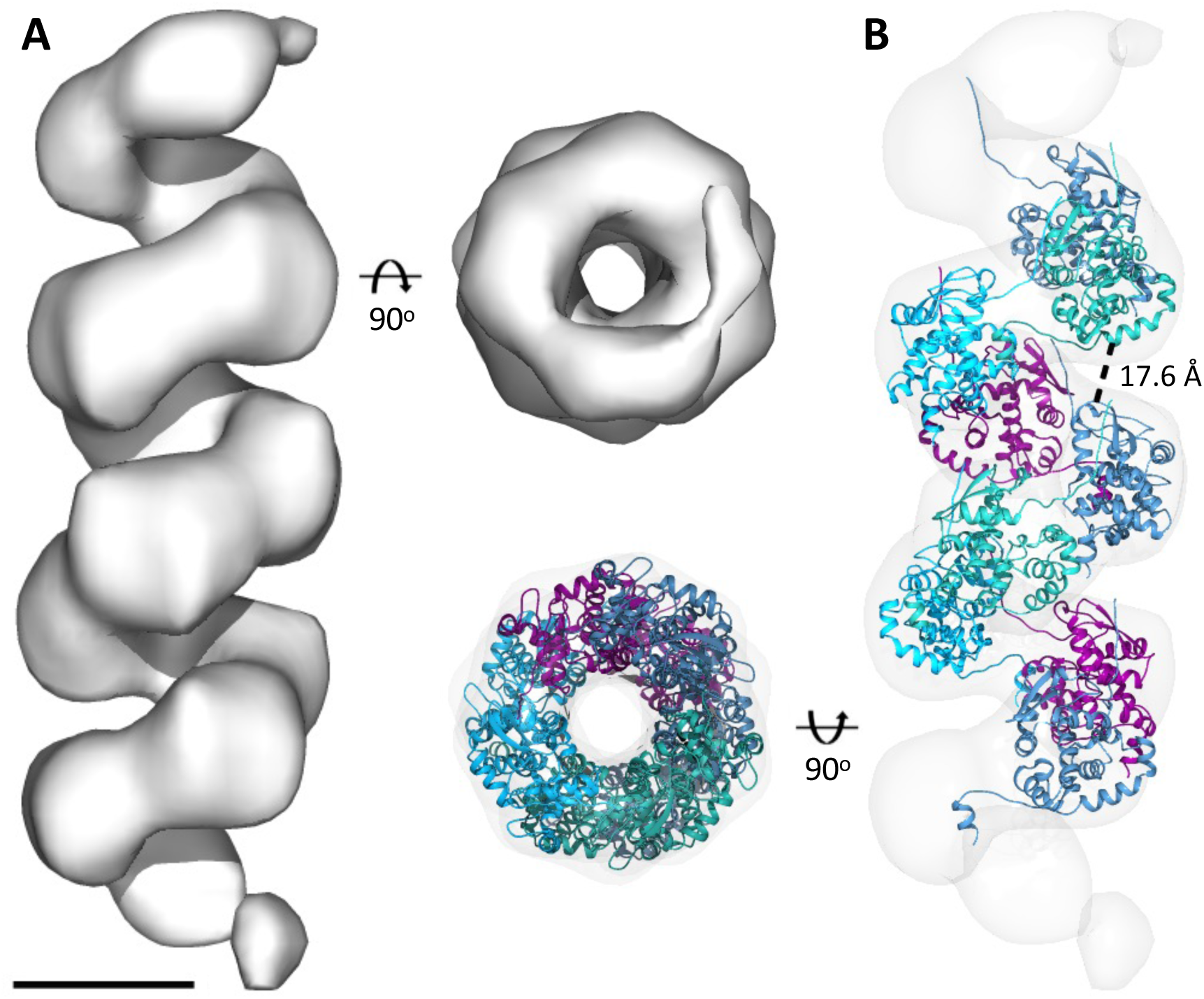
Cryo-EM model of the BUNV RNP. **(A)** Cryo-EM average with helical symmetry generated a 13 Å model of a BUNV RNP filament. (**B**) Flexible fitting a ‘split’ NP model with appropriate restraints produced a pseudo-atomic model of the BUNV RNP. Distance between rungs of the helix indicated. Scale bar: 5 nm.

Given that helical RNPs have 3.7 NP per turn, 11 bases per NP and a 6.3 nm pitch, the lengths of S, M and L segments (nucleotides per segment/(3.7*11))*6.3) were predicted as 149, 690 and 1064 nm, respectively, very close to the measured mean lengths of 160, 650 and 1050 nm (Figure S7A). This calculation validates the correct identification of the S, M and L segments, that they are helical and that we have obtained the correct helical parameters.

### NP crystal structures and molecular dynamics allow the generation of an atomic model for the BUNV RNP

The generation of a 3D helical model of a BUNV RNP permitted the fitting of a ‘split’ NP crystal structure (Figure S13) to generate a pseudo-atomic model. In all available crystal structures of multimeric orthobunyavirus NP the globular core of the protein is highly conserved (Figure S14A), as are the binding positions of the N- and C-terminal arms relative to their bound neighbour (Figure S14B). To preserve these aspects of the structure we produced a model of what we have termed ‘split’ NP (Figure S13C) that contains the core of a single NP monomer and the bound arms of its two neighbours. A limitation of this approach is that it prevented us from modelling the viral RNA along the NP arms, and therefore we did not include it in our model. The NP structure was symmetrized and molecular dynamics was applied restraining the ‘split’ NP. This allowed freedom of movement of the linker region of each terminal arm, which has already been shown to be flexible, while minimising alterations to the globular core of the protein or to the binding regions of the arms (Figure S14A). As a result, we obtained a helical NP model for BUNV RNP (Figures 3B and 4A). Displaying the surface coulombic potential of the helical model confirms that the positively-charged RNA-binding groove of each monomer is aligned in a way that maintains an RNA-binding channel along the interior of the entire helical filament (Figure S15A). Studying the surface coulombic potential also ruled out the possibility of longitudinal interactions between charged areas on the surface of monomers, as the surfaces of the protein that face each other longitudinally contain mainly negatively charged residues (Figure S15B). Furthermore, these rungs are too far apart (approximately 18 Å) to permit intermolecular interactions of any kind between NP monomers along the longitudinal axis (Figure 3B).

**Figure 4.**
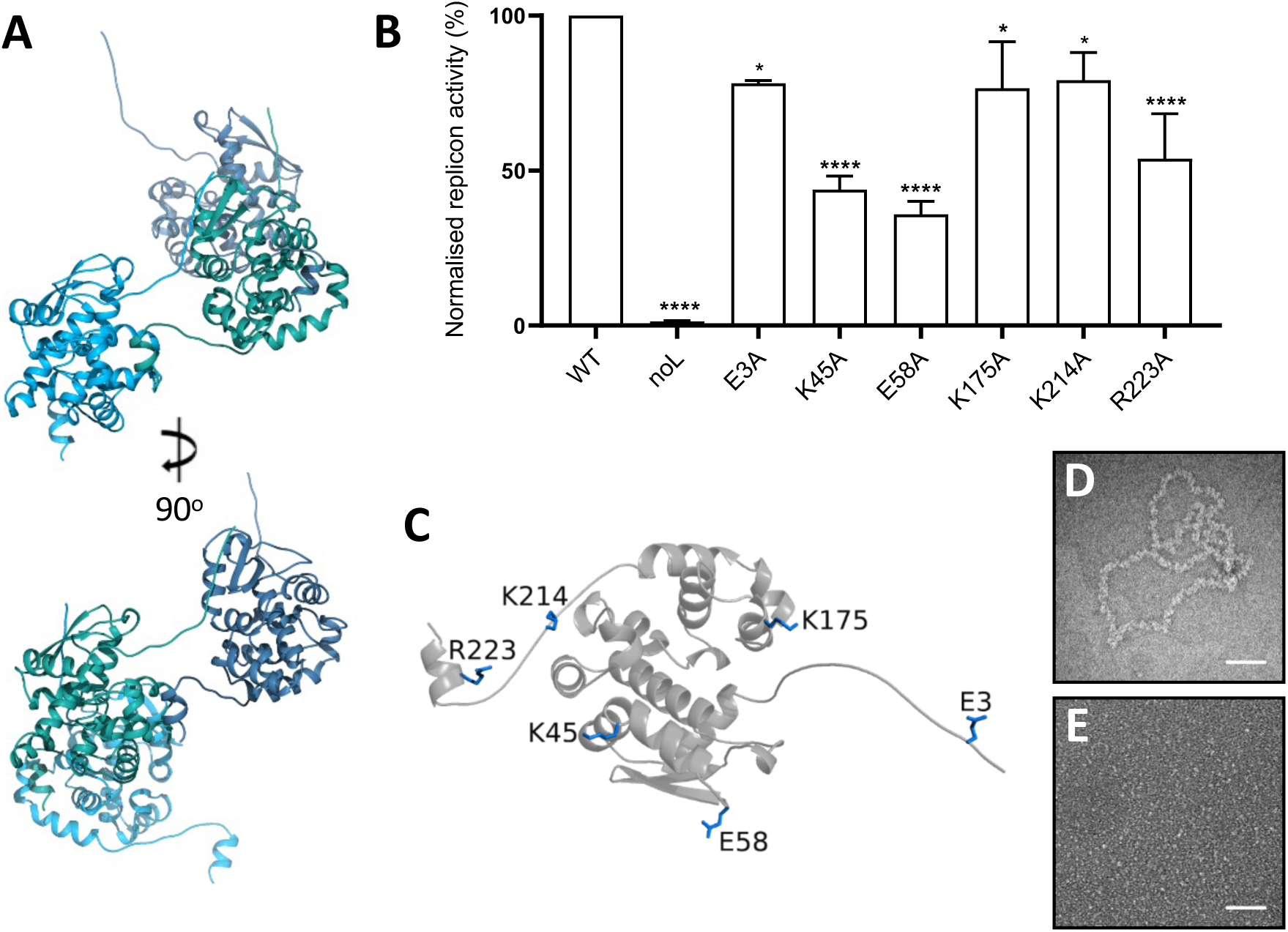
The BUNV RNP helix is maintained by the RNA. **(A)** Orientation of NP within the BUNV RNP model. (**B**) Histogram showing gene expression activity of mini-genome RNPs reconstituted using WT and mutant NPs, alongside transfections in which the BUNV polymerase was omitted (noL). *: p ≤ 0.05, **: p ≤0.01, ***: p≤0.001, ****: p≤0.0001. (**C**) Tested mutants highlighted on an NP monomer from the RNP helical model. (**D**) Untreated BUNV RNPs and (**E**) BUNV RNPs treated with 20 μg of RNase A, visualized using negative staining EM. Scale bars: 50 nm.

### The contribution of RNA to maintaining the helical RNP architecture

The generation of our helical NP model allowed investigation of the mechanism of native RNP assembly, in terms of formation of both the NP-NP multimer chain, and its helical architecture. The comparison of our crystallographic model of the closed-ring BUNV NP tetramer with that of the native helical RNP allowed the identification of 7 additional residues with sidechains potentially involved in weak H-bond or salt bridge NP-NP interactions within the RNP (Supplementary Tables S1-S3). As the resolution of our map prevented us from refining the side chains of NP, to test their involvement in helical RNP formation, each of these 7 residues were individually substituted for alanine in reconstituted RNPs, and their resulting function tested by mini-genome analysis, using reporter gene expression as a measurement of active RNP assembly (Figures 4B and C). First, we tested if the mutations perturbed NP expression. All mutants tested except E173A resulted in a level of expression comparable to that of wild-type NP (Figure S16). We then tested the mutations that did not affect NP expression using a mini-genome assay, and all of these resulted in a significant reduction of mini-genome reporter activity (Figure 4B). This finding establishes important roles for these residues in the formation of an active RNP and thus further validated our helical NP model. The location of all these residues at the interface between laterally-adjacent NP monomers, and the distance between the helix rungs suggests they are solely involved in lateral NP multimerization, although, we cannot rule out the possibility that they contribute to RNP helical architecture by imposing torsion within the NP chain. They could also be involved in RNA binding and/or RNP packaging. Additionally, the fact that none of the mutations completely abolishes the expression of the mini-genome system, suggests several residues might be involved in helical NP-NP interactions.

However, by a process of elimination, our RNP model implicates the only other component of the RNP, namely the RNA, as a possible contributor to maintaining the helical RNP architecture. To test this possibility, purified BUNV RNPs were incubated with RNase A at concentrations known to degrade BUNV NP-encapsidated RNA (35), and then visualized by negative staining EM. Characteristic BUNV RNPs were abundant in untreated samples (Figures 4D and S17A), whereas following RNase treatment, no RNPs were detected (Figures 4E, S17B and C). Instead, a diffuse background staining was apparent, most likely comprising dissociated NP monomers, not observed in untreated samples. Of note, intact NP monomers were present in all these samples, as assessed by western blot (Figure S17D), suggesting NP-NP dissociation and not protein degradation was responsible for the loss of intact RNPs. Taken together, these findings suggest that both the encapsidated RNA and the NP-NP lateral interactions are important for the integrity of the flexible orthobunyaviral RNP.

## Discussion

We present the first 3D model of a native orthobunyaviral RNP, derived using a combination of EM approaches and molecular dynamics modelling. The model shows that the BUNV RNP exhibits clear helical architecture, and reveals key information about its assembly, structure and function.

Our current and previous EM observations show that the orthobunyavirus RNP exhibits a high degree of flexibility and thus heterogeneity, which has hampered the application of single particle techniques to derive high-resolution structural models of RNPs from this group of viruses. To overcome this hurdle, here we combined different EM approaches. Negative staining allowed us to conclude that the width and length of the RNPs, and the distances between NPs, were compatible with a helical RNP. AFM allowed the determination of the helical handedness. Sub-tomogram averaging and cryo-EM single particle using a cylinder as an initial reference further confirmed the helical arrangement of the RNP; however, the flexibility of the RNP prevented the selection of straight segments, and application of computational helical reconstruction techniques. Therefore, we followed a single particle approach using the helical sub-tomogram average as a reference, and only at the last stages of the process searched for and applied helical symmetry. The resulting average at 13 Å, combined with restraints derived from crystallographic analysis of deposited orthobunyavirus NP oligomers, allowed us to apply molecular dynamics to obtain a helical NP model for BUNV RNPs. The buffer in which our RNPs were visualised was selected for the ease of finding straight sections of filament to analyse, and we acknowledge that this is not necessarily physiologically relevant. However, during the testing of a range of buffers the overall architecture of the RNPs did not change (Figure S6).

As suggested by crystallographic models of orthobunyavirus NP tetramers with bound RNA, our helical BUNV RNP model shows the genomic RNA to be sequestered on the inner face of the helix, where it lies within an RNA-binding cleft that forms a continuous tract within the rising helical arrangement of NP monomers. The NP oligomerisation arms also appear to partially shield the RNA from the solvent at lateral NP-NP interfaces, as was observed in an NP crystal structure from SBV (15) and together this encapsidation strategy would prevent the RNA from extruding out of the RNP helix. This has significant ramifications in relation to the bunyaviral replication cycle. For instance, it seemingly rules out RNA-RNA intersegment interactions that may contribute towards a selective orthobunyavirus segment packaging mechanism, in contrast with influenza virus, for which extensive inter-segment interactions are proposed to drive selective packaging of all 8 genome segments during assembly (40). The almost complete coverage of BUNV RNA within the RNP is consistent with the current model for *Bunyavirales* segment packaging, for which no evidence of segment selection has been found (41).

The flexibility of BUNV RNPs contrasts sharply with the more rigid RNPs belonging to members of the *Mononegavirales* order of non-segmented negative-sense RNA viruses, exemplified by measles and Ebola viruses (42–46), or the more closely-related *Orthomyxoviridae* family of segmented negative-stranded RNA viruses, such as influenza virus (39, 47, 48). Our 3D model shows the molecular basis for its flexibility lies in the lack of NP-NP interactions along the longitudinal RNP axis, dictated by the distance between NP monomers on adjacent rungs along the helix. Instead, the NP-NP chain in the RNP helix is maintained by lateral interactions involving the N- and C-terminal arms (11–16), as well as additional residues that we identify here, which engage in weak H-bond or salt bridge interactions. Furthermore, our results suggest the maintenance of the architecture of the RNP is dependent on the presence of encapsidated RNA.

The level of flexibility observed in the native helical BUNV RNPs permits a large degree of bending (up to 90°; e.g. Figure 1A) while still apparently preserving the integrity of the filament, which is likely of great benefit for several stages of the replication cycle. Flexibility will facilitate segment packaging during virion assembly, as it will allow the genomic segments to contort within the virion interior. BUNV particles are pleiomorphic, with diameters of around 108 nm (10), and so BUNV S, M and L segments, with lengths of 150, 650 and 1050 nm, respectively, clearly could not be packaged if their RNPs were rigid rod-like structures. Interestingly, the virion diameter of 108 nm suggests an internal volume of around 6.7 × 10^5^ nm^3^, and by treating the three BUNV RNPs as a single cylindrical volume with a radius of 4.6 nm (Figure 1B) and combined length of 1850 nm, the volume that the three segments would occupy according to our model is 1.2 × 10^5^ nm^3^, approximately one fifth of the capacity of the virion. Thus, it is entirely plausible that BUNV segments are packaged within virions without the need for further condensation.

RNP flexibility also allows the BUNV segments to adopt their characteristic circular RNP conformation in which their 3’ and 5’ ends interact, and this ability is particularly important for the S segment, for which circularization would require the shortest radius of curvature due to its short nucleotide length. Segment circularization is important for BUNV gene expression, and appears critical for recognition of the RNPs by the RdRp (32, 49), thus RNP flexibility is a critical requirement for virus viability.

The current model for how bunyaviral RNPs circularize is unclear, with possible contributions from inter-terminal Watson-Crick base-pairing interactions, and interactions with sequence-specific binding sites on the RdRp surface. Our negative staining EM observations revealed BUNV RNPs to be almost exclusively circular molecules, with less than 1% appearing in a linear form. Interestingly, close inspection of the circular RNPs revealed no evidence of where the joins between the RNA 3’ and 5’ termini might be. This finding was unexpected, as the inter-terminal interactions shown to be important for BUNV RNA synthesis (50–52) would be predicted to result in the formation of a partial duplex by Watson-Crick pairings, distinct from all other segment sequences that are single stranded. As the NP RNA-binding groove cannot accommodate double stranded RNA (11–15), this terminal hairpin would be expected to be displaced from the oligomeric NP chain, resulting in a region of the RNP with a distinctive structure. We were unable to detect any such regions, although this may reflect the relatively low resolution of negative staining EM.

An alternative mechanism for segment circularization is that it is mediated by the RdRp, based on the observation that the 3’ and 5’ terminal sequences bind to separate sites on the RdRp surface (32, 49). However, most of the RNPs we imaged lacked a putative RdRp, suggesting that RNP circularization is not dependent on a resident RdRp to tether their segment termini. Circularization independent of the presence of an RdRp raises the possibility that interactions between adjacent NP molecules alone may drive segment circularization, leading to the formation of a continuous NP chain without distinctive ends; however, complementary ends are critical for the viral life cycle therefore the genome ends must interact at some point in the infectious cycle. It is of note that native RdRp has never been observed in association with orthobunyaviral RNPs, in contrast to that of influenza virus for which the RdRp readily co-purifies as part of the RNP complex (39, 48). This might suggest weak RdRp interactions within bunyavirus RNPs.

Our RNP model (Figure 3) shares some similarities with the previously described helical array of crystallised LACV NP (12) such as the helical pitch and the general arrangement of the NP molecules. However, the crystal structure has a regular four monomers per turn of the helix and very closely resembles the tetrameric crystal structure, while our model has 3.7 monomers per turn. Therefore, the crystal structure could be affected by crystallographic packing artefacts. We also note that the crystal structure is derived from apo NP, and this may have important structural consequences, as our results suggest that the encapsidated RNA plays an important role in helix maintenance. Interestingly, the recently solved structure of an *in vitro* reconstituted Hantaan virus (HTNV) RNP is similar to our model in terms of its helical pitch and number of subunits per turn (Figure S18A) (22). However, in contrast to our model, the HTNV RNP model shows a clear role for longitudinal NP-NP interactions in helix maintenance, with each NP monomer contacting six others across multiple rungs of the helix. This results in a structurally-homogeneous, rod-like RNP distinct from the highly flexible native BUNV RNP we characterise here. Many of the residues within the HTNV NP which the authors ascribe to these longitudinal NP-NP interactions (22) have no equivalent in the NP of BUNV, which is structurally distinct (Figure S18). It is likely that this marked difference in the capacity to form NP-NP interactions leads to differences in NP oligomerisation, which in turn result in the very different RNP structures of the two viruses. This finding, along with significant differences in NP structure and multimerization strategies across the five animal and human infecting families that make up the *Bunyavirales* order suggests their corresponding RNPs may also show significant differences in overall architecture, with potential implications for the design of therapeutics that target this order of viruses.

## Data Availability

The asymmetrical and symmetrical averages and pseudo-atomic model are deposited under EMDB accession numbers EMD-11847 and EMD-11849, and PDB accession number 7AOY.

## Acknowledgements

We thank the Astbury Biostructure Laboratory (ABSL) Facility Staff, especially Martin Fuller and Rebecca Thompson, for assisting with negative stain and cryo-EM data collection, and Neil Ranson for securing funds.

## Supplementary Materials and Methods

### Virus Propagation and Purification

Baby hamster kidney (BHK-21) cells at 80 – 90% confluency were infected with BUNV at a multiplicity of infection (MOI) of 0.01 and maintained in serum free DMEM at 32°C. After 96 hours supernatant was collected and clarified by centrifugation at 3700 *g* for 20 minutes. Virus particles were then purified and concentrated by centrifuging at 100000 *g* for three hours in an SW32 rotor (Beckman-Coulter) with an underlay of 30% D-sucrose prepared in TNE buffer (100 mM Tris-HCl, 200 mM NaCl, 0.1 mM EDTA, pH 7.4) supplemented with protease inhibitors (Roche). Pellets were resuspended in 100 μl of TNE buffer at 4°C overnight and pooled together the following day.

### RNP Purification

Virions were disrupted by a single freeze/thaw cycle, 30 minute incubation in 1% Triton X-100 (Sigma-Aldrich) and 0.1% NP-40 alternative (Calbiochem) or 30 minute incubation in 30% D-sucrose in TNE buffer. Disrupted virions were applied to the top of a 10-25% OptiPrep gradient prepared in TNE buffer and centrifuged in an SW60 rotor (Beckman-Coulter) at 250000 *g* for 90 minutes. Fractions were collected and analysed for the presence of BUNV NP by western blotting using specific antisera (produced in-house), and those with the strongest NP signal were buffer-exchanged to remove OptiPrep. A range of buffers was tested (Figure S6) and a reduced-salt variant of TNE containing only 25 mM NaCl was used for all further analysis by EM.

### Negative-staining Electron Microscopy

Immediately following buffer-exchange, 5 µL aliquots of RNPs were adsorbed to glow-discharged, carbon-coated copper grids for three minutes. Following two water washes, grids were stained for 30 seconds with 1% uranyl acetate. Images were collected on a 120 kV FEI Tecnai G^2^-Spirit microscope at a nominal magnification of 45000X.

RNPs were subject to autopicking in Relion 3.0 (1) and a total of 175,583 particles were extracted. Rounds of 2D classification were used to discard the most heterogeneous particles and a final set of 14,899 particles was used to generate 2D class averages. To estimate the RNP length, 609 discrete RNP filaments were manually traced in Fiji (2) using the line tool and the length of the line was determined and used as an estimation of RNP length. In a similar manner, the line tool was also used to measure filament diameters and the distance between neighbouring subunits from within class averages.

### Cryo-electron Microscopy

Carbon-coated quantifoil grids were prepared by plunge-freezing in liquid ethane using an FEI Vitrobot IV. Cryo-ET tilt-series were collected on an FEI Titan Krios microscope operating at 300 kV and equipped with an energy filtered Gatan K2 XP Summit and Volta phase plate. Tilt-series were collected at 2° intervals from -60° to 60° at a nominal magnification of 71000X for a pixel size of 2.7 Å. During tilt series and single particle data collection, on-the-fly motion correction and CTF estimation were set up in Relion 3.0 as previously described (3).

Tomograms were reconstructed within the eTomo pipeline for fiducial-less alignment using weighted back projection from the IMOD software package (4). Sub-tomogram averaging (STA) was carried out with and within PEET (5). Sub-tomograms were manually picked using 3dmod and STA was performed by alignment and averaging of these sub-volumes in PEET, using absolute value of cross-correlation and strict search limit checking. For the first iterations only the orientation of the RNP (without rotation) was included in the angular search range, starting with a maximum of 24° and a step of 8° and halving with each iteration, to produce a straight cylindrical model. In later iterations when the RNPs were orientated down to 6°, the rotation search was included.

Single particle data was collected on an FEI Titan Krios microscope operating at 300 kV and equipped with a Falcon 3EC. Micrographs were collected at a nominal magnification of 130000X for a pixel size of 1.07 Å.

Data were processed in Relion 3.0 (1). Approximately 1000 particles were picked for reference-free classification and the resulting classes were used as templates for autopicking. 1.47 million particles were selected by autopicking and extracted with a box size of 274 Å, binning particles 4 times (resulting in pixel size 4.28 Å). Two rounds of 2D classification selected the most homogenous 135102 particles. To validate the helical nature of the RNP, these particles were first subjected to 3D classification using a cylinder as a reference (removing any possible bias towards a helical model). The resulting averages were clearly helical, and therefore the particles were taken forwards for image processing using the STA-derived model. The selected particles were 3D classified against the STA model and two further rounds of 3D classification left the 5077 most homogenous particles. These particles were re-extracted with a smaller box size of 214 Å to increase homogeneity and were subject to two more rounds of 3D classification. In the final round of 3D classification the particles were divided into classes of approximately 4000 particles, as was recently done for IAV (6), and the top class was subject to helical reconstruction without symmetry. The same 5077 particles were then re-extracted with the increased box size of 274 Å to aid in alignment and a final refinement was carried out with helical symmetry applied. Local searches were applied that centred on values of 100° for twist and 15 Å for rise, as determined by viewing an XYZ projection of the non-symmetrical model. These always converged on the same values of -104.9° for twist and 17.7 Å for rise so these numbers were used for searches for the final refinement with helical symmetry imposed, which converged on -105.26° twist and 18.25 Å rise. The asymmetric and symmetric reconstructions are highly similar (their cross-correlation value is 0.94, when using chimera fit in map tool), suggesting we found the correct helical parameters.

### Atomic modelling of BUNV RNPs

To generate an atomic model of BUNV RNPs, we used the coordinates of BUNV NP as a starting point (PDB: 3ZLA (7)). First, a trimer of NP was rigid-body fitted into the cryo-EM average using Chimera (8). Of the two possible orientations, one of them (which we term ‘beta-strand up’) fit slightly better into the non-helically symmetrised average (it had an average map value at fit atom positions of 0.09 vs. 0.081 for the alternative orientation); however, both orientations were taken forward to allow comparison of the final result. As has been previously reported, the core domain of all atomic coordinates from orthobunyavirus NP is constant, while the N- and C-terminal arms are flexible (Figure S14 and (9)). However, when comparing the NP core with the arms of the adjacent subunits (which we term ‘split’ NP), the interactions between the arms and the core domain remain constant across the structures of all orthobunyavirus NP oligomers (Figure S14). Therefore, we next fitted a ‘split’ NP into the central monomer of the rigid-body fitted trimers, and manually added the missing amino acids of the coordinates of BUNV NP in COOT (10) (corresponding to the linker regions between the arms and the core domain), to prevent the movement of the terminal ends of the protein to positions that are not physically possible during flexible-fitting. The resulting NP monomer was helically symmetrised in Chimera, and the NP helix was flexible-fitted into the cryo-EM average using MDFF (11), following the program’s guidelines. The ‘split’ NP domains were kept as rigid domains by applying domain restraints, and symmetry restraints were also employed. Of the two possible orientations of the NP, the ‘beta-strand up’ resulted in a better model, in terms of positive electrostatic charges for interactions with the RNA.

### PISA

PDBePISA (Protein Interfaces, Surfaces, and Assemblies: http://pdbe.org/pisa/) and Chimera (8) were used to predict inter-chain interface contact residues, including H-bonds and salt-bridges, involved in the interaction between BUNV NPs within the previously published BUNV NP tetramer (PDB: 3ZLA (7)) and within the helical RNP model.

### BUNV Replicon Plasmids and Mutagenesis

Previously described plasmids were used for the BUNV replicon system assay (12). Briefly, the replicon system is constituted by BUNV L and NP support plasmids (pT7riboBUNL, pT7riboBUNN) which express BUNV L and NP downstream of the T7 pol promoter and the internal ribosome entry site (IRES) of encephalomyocarditis virus (ECMV), the minigenome pT7riboBUN-SREN, which contains the Renilla luciferase gene in the negative sense flanked by the BUNV S segment UTRs, and the control plasmid pCMV-Firefly-Luc, which expresses firefly luciferase under a cytomegalovirus (CMV) promoter. Mutagenesis of the NP open reading frame (ORF) was achieved using pT7riboBUNN plasmid as template for site-directed mutagenesis (SDM). SDM primers were designed using NEBaseChanger and SDM reactions were performed using a Q5 SDM Kit (NEB) according to the manufacturer’s instructions. Corresponding DNA clones were transformed into competent bacterial cells, purified using Miniprep kits (Qiagen) and mutations were confirmed using Sanger sequencing.

### BUNV Replicon Transfection and Luciferase Activity Assay

BSR-T7 cells were seeded in 24-well plates 1 day prior to transfection at a density of 5×10^4^ cells per well. Cells were transfected with 0.2 μg each of pT7riboBUNL, pT7riboBUNN or one of the mutated NP clones, the minireplicon plasmid pT7riboBUN-SREN and 0.1 μg pCMV-Firefly-Luc (internal transfection control). TransIT-LT1 Transfection Reagent (Mirus Bio) was used as transfection reagent in a 2.5:1 ratio of TransIT-LT1 (μl):DNA (μg) according to the manufacturer’s instructions. At 24 h post-transfection, Renilla and firefly luciferase activities were measured using the dual-luciferase assay kit (Promega). Statistical analyses were performed using one-way ANOVA, followed by Dunnett’s multiple comparisons test to determine significant differences in replicon activity between NP mutants and WT BUNV NP.

### AFM sample preparation and imaging

The handedness of fibrils was determined using AFM. 18 µL of RNPs was deposited on freshly cleaved mica in an imaging buffer of 10mM NiCl_2_, 10 mM HEPES pH 7.5. All AFM imaging was carried out at room temperature with the RNPs in solution and hydrated in the imaging buffer. AFM observations were performed in tapping mode using a Dimension FastScan Bio with FastScan-D-SS probes (Bruker). The force applied by the tip on the sample was minimized by maximizing the set point whilst maintaining tracking of the surface. Images were processed using first-order line-by-line flattening to remove the sample tilt and background using Nanoscope Analysis (Bruker). Correlation averaging was performed using self-written code in MATLAB. RNPs were first skeletonized to trace the profile before profile smoothing and digitally straightening. A reference section within the digitally straighten image containing 1 full helical repeat of the RNP was then cross correlated along the entire length of protein to search for similar features. Correlation averages were created using the 80 of 120 sections with the highest cross correlation coefficient. Cross-correlation was also performed with a vertically flipped reference image to search for possible left-handedness. The code employed for AFM image analysis has been deposited in https://github.com/George-R-Heath/Correlate-Filaments.

---

**Figure S1.**
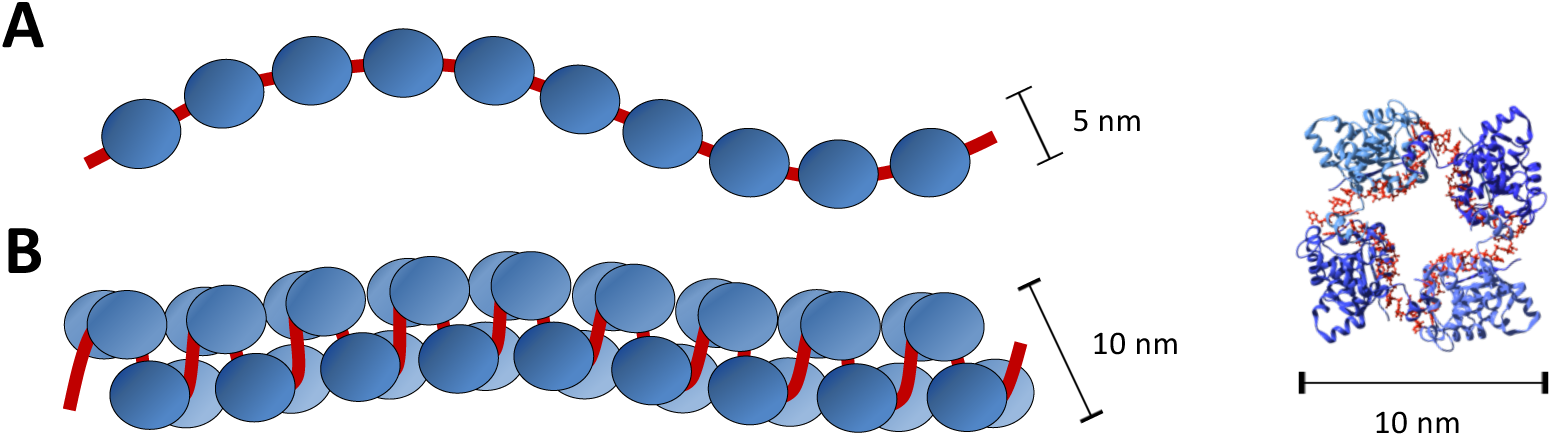
Proposed models of NP organisation within orthobunyavirus RNP. **(A)** monomer-based ‘beads on a string’ and **(B)** tetramer-based helix alongside the crystal structure of tetrameric BUNV NP.

**Figure S2.**
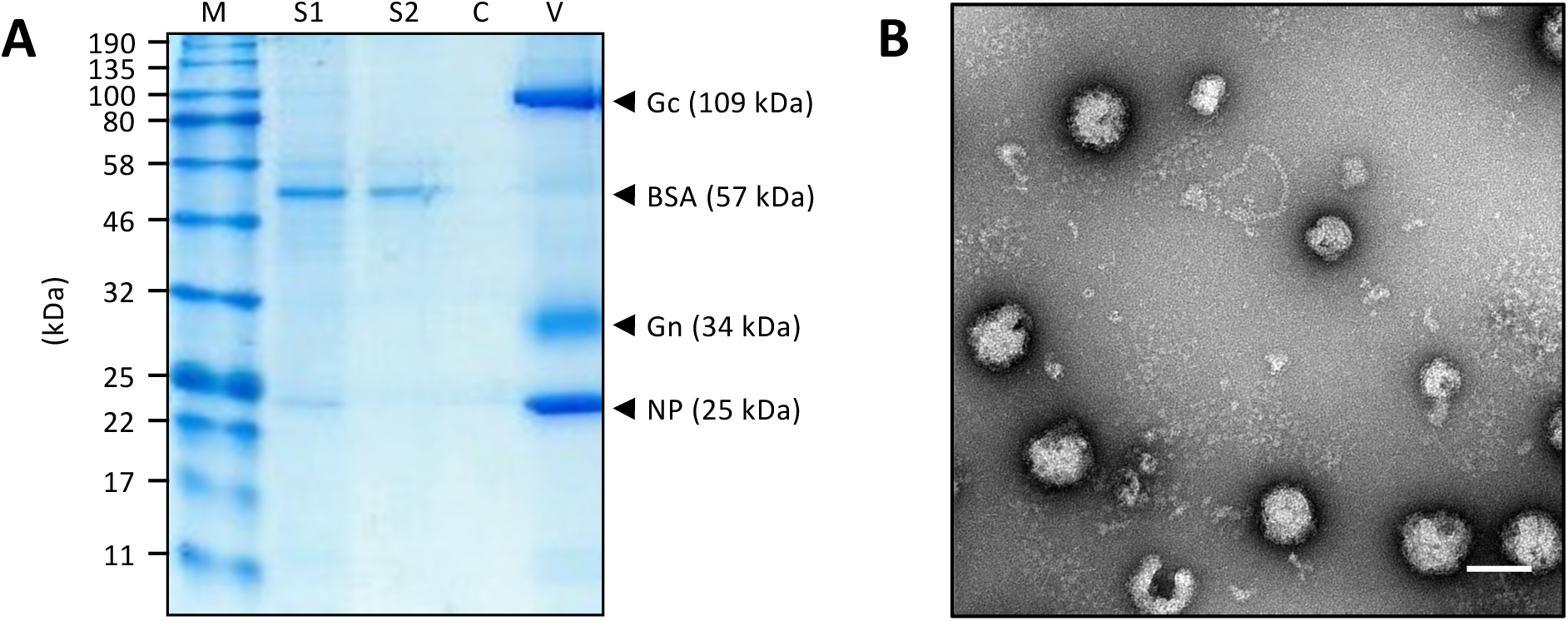
Purification of BUNV. (**A**) SDS-PAGE analysis showed an increased concentration of virus after pelleting through a sucrose cushion, with high purity and removal of BSA contaminants. M = marker, S1 = supernatant pre-purification, S2 = clarified supernatant pre-purification, C = cushion post-purification, V = resuspended viral pellet. (**B**) Negative staining EM confirmed purity and abundance of virions. Scale bar: 100 nm.

**Figure S3.**
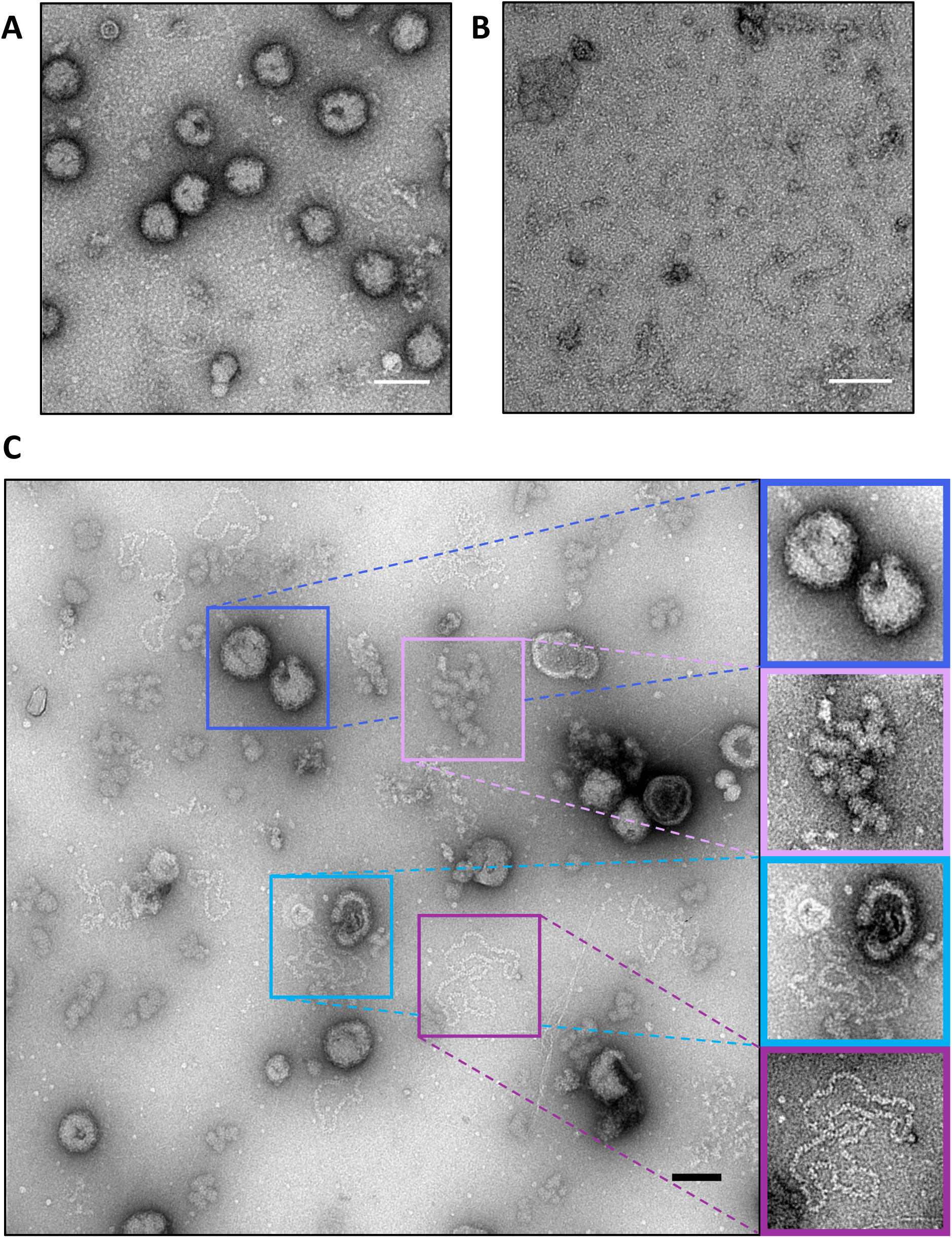
Methods of BUNV RNP release, examined by negative staining. (**A**) Sucrose washes were not reproducible and often failed to lyse virions. (**B**) Washing with detergent was effective at disrupting virions but produced a high level of background signal. (**C**) Freeze/thaw was highly effective for release of viral RNPs. From top to bottom, insets show mostly intact virus, debris attributed to disrupted viral membranes, RNPs spilling from their parent virus and discrete RNPs released from virions. Scale bars: 100 nm.

**Figure S4.**
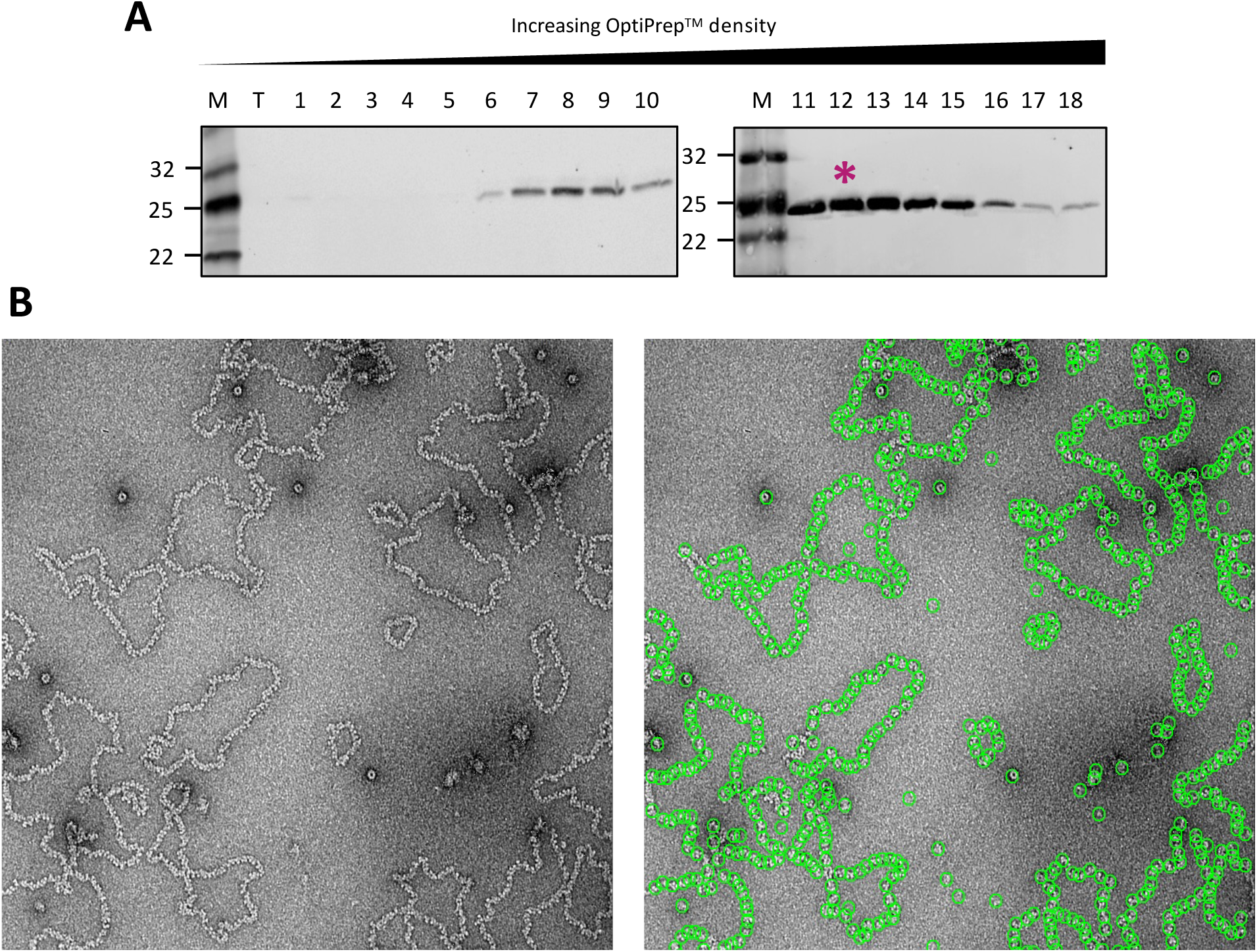
BUNV RNP purification. (**A**) Fractionation and western blotting showed the location of RNPs within a continuous density gradient (asterisk). (**B**) Representative micrograph showing abundant, purified BUNV RNPs (left), and the effectiveness of the Relion autopicking tool for selecting particles (right).

**Figure S5.**
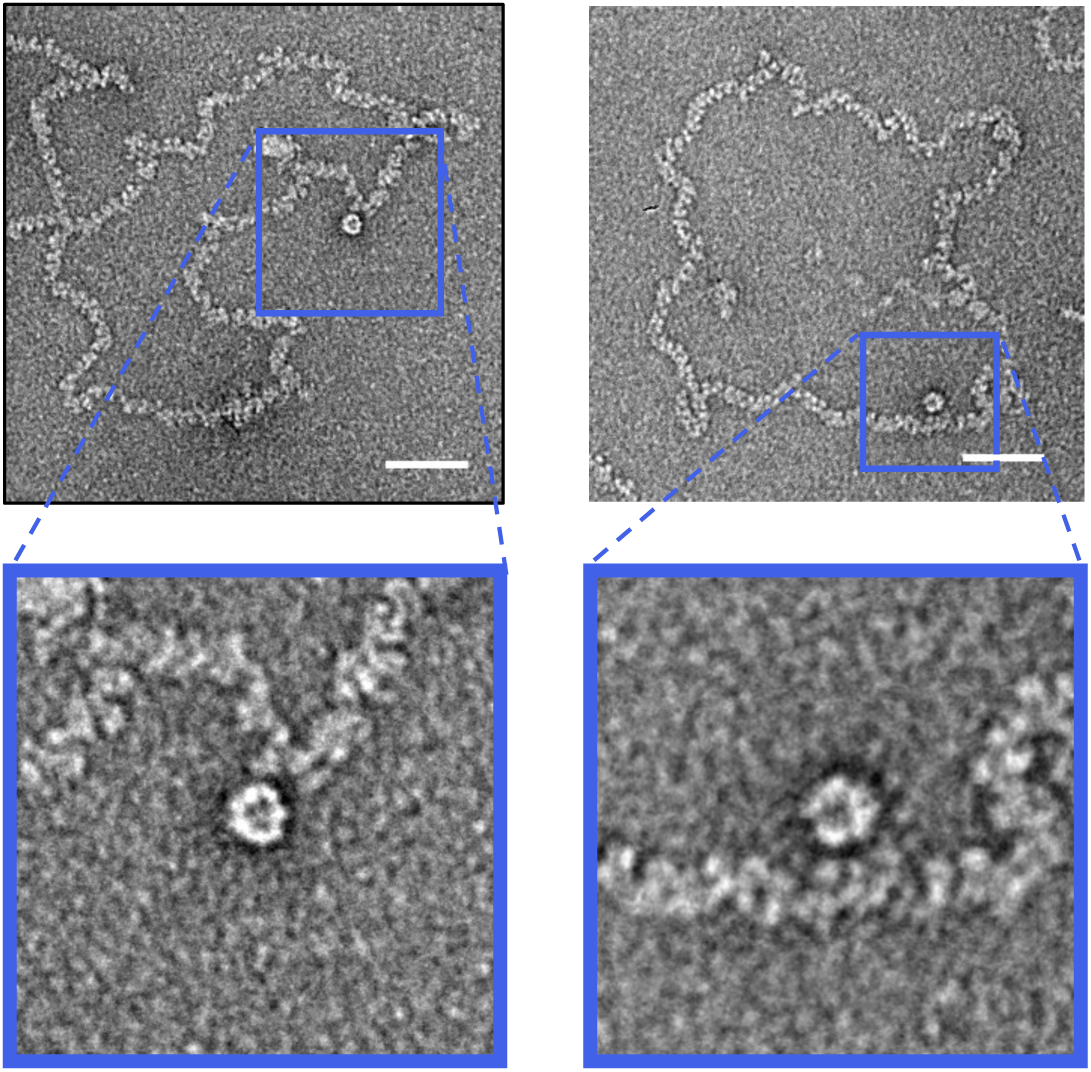
Putative polymerase molecules within BUNV RNP preparations. During analysis of negatively stained BUNV RNPs, a population of large structures with resemblance to viral polymerases was found, often in association with RNP filaments. Scale bars: 50 nm.

**Figure S6.**
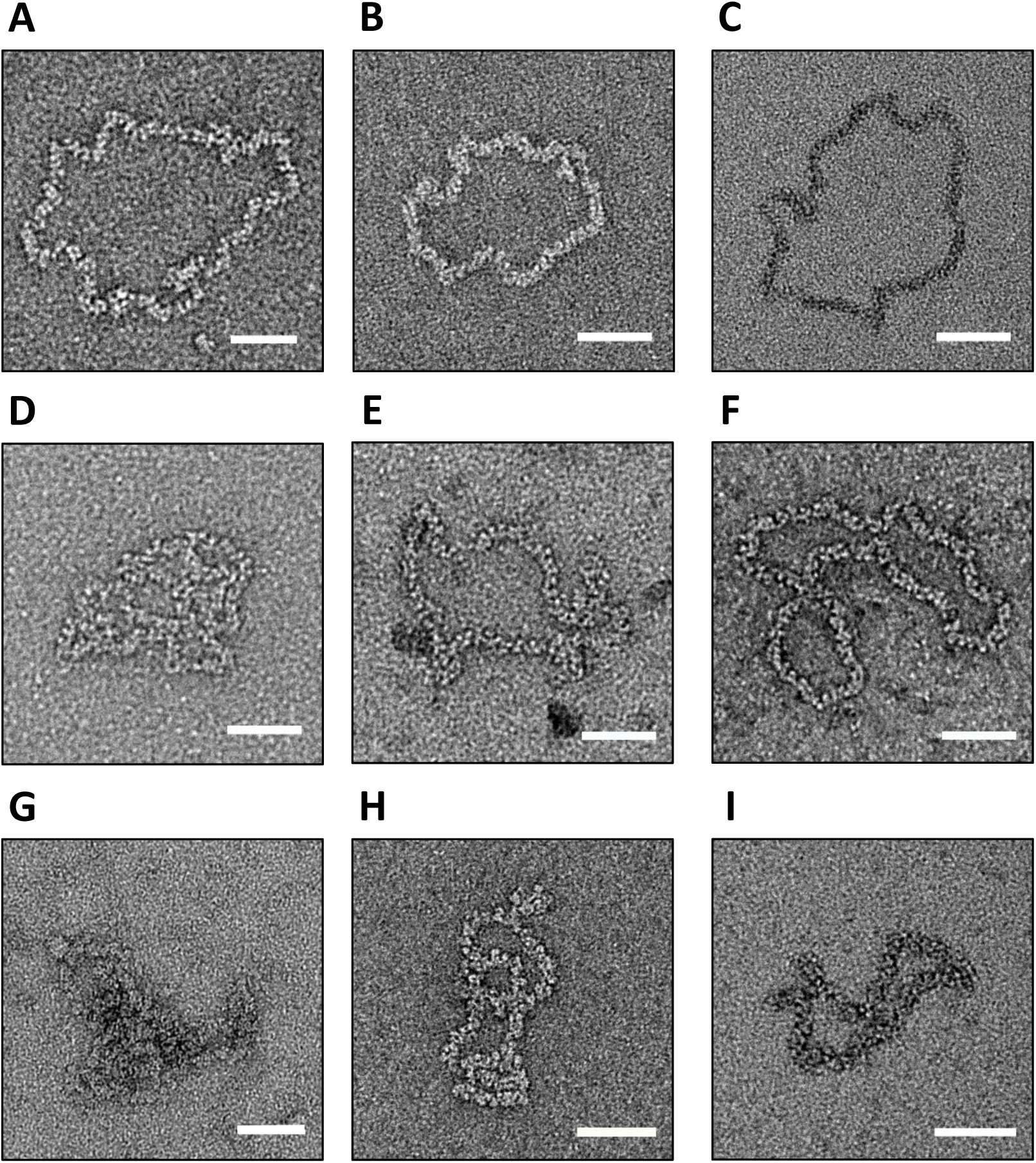
Visualization of purified BUNV RNPs after buffer exchange into different conditions. (**A**) TNE with no salt. (**B**) TNE with 25 mM NaCl. (**C**) TNE with 50 mM NaCl. (**D**) TNE with addition of MgCl_2_. (**E**) TNE with addition of DTT. (**F**) TNE with KCl substituted for NaCl. (**G**) TNE with addition of antisera. (**H**) TNE pH 6.4. (**I**) TNE pH 8.0. Scale bars: 50 nm.

**Figure S7.**
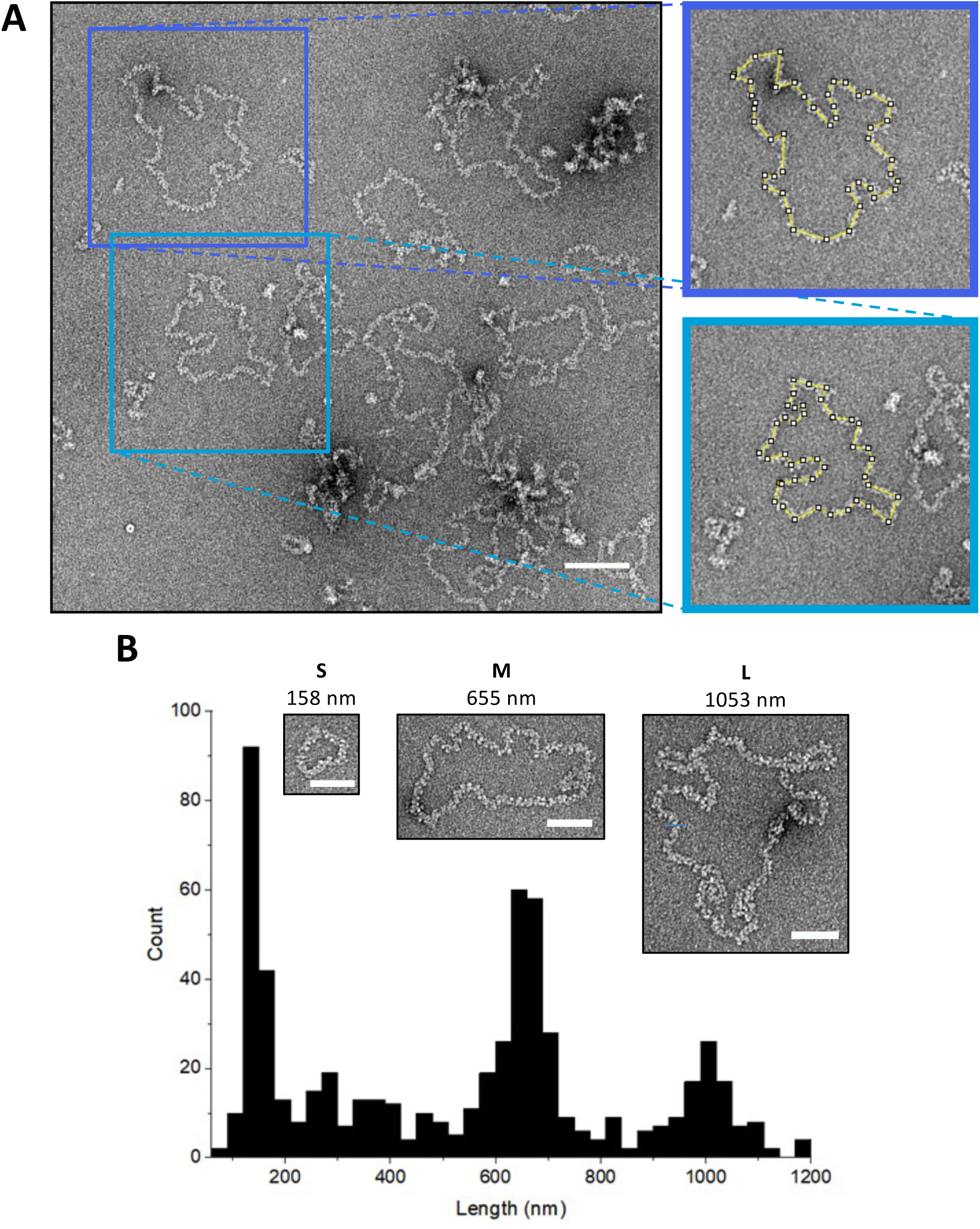
BUNV RNP length measurements. (**A**) Schematic of discrete RNP tracing. Scale bar: 100 nm. (**B**) Tracing the lengths of filaments revealed three distinct populations corresponding to S, M and L genomic segments. Insets show representative S, M and L RNPs and their lengths. Scale bars: 50 nm

**Figure S8.**
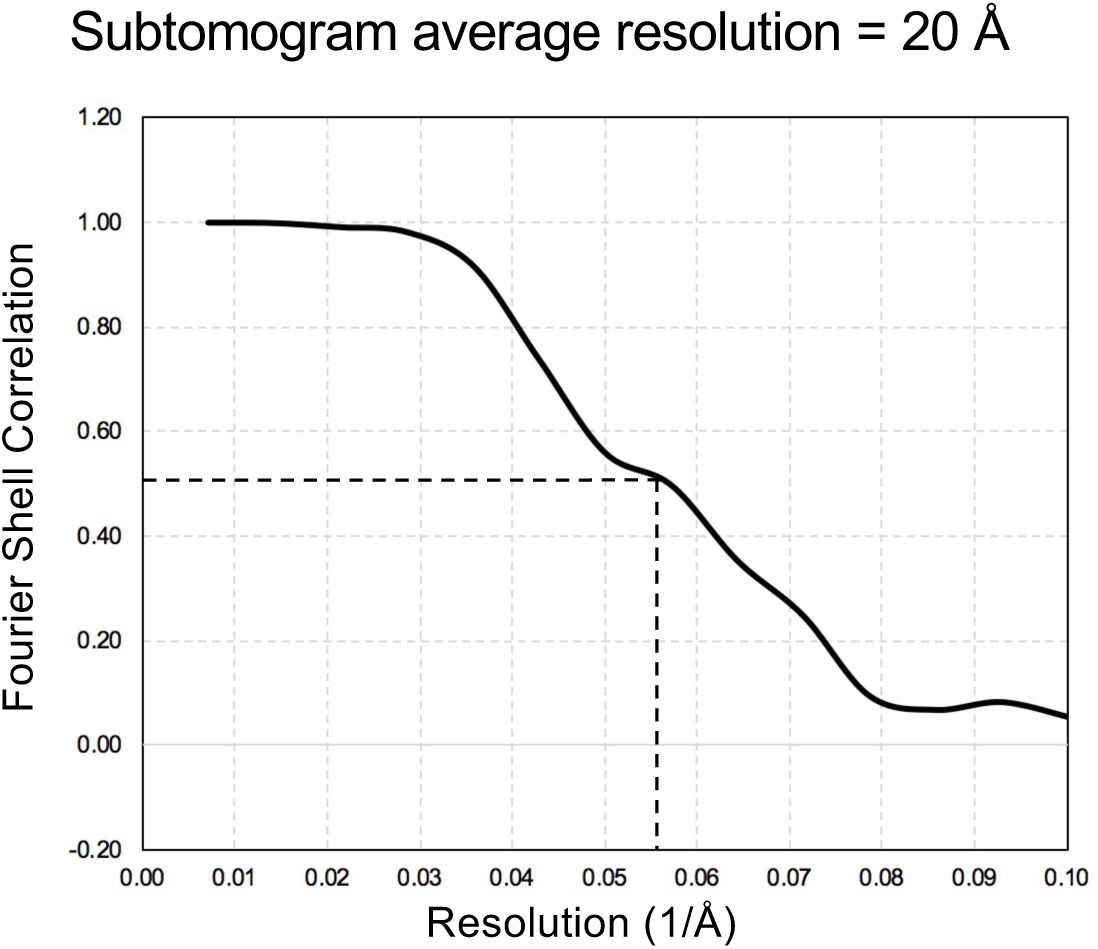
FSC curve of the sub-tomogram average. Resolution is 20 Å at an FSC cut-off 0.5.

**Figure S9.**
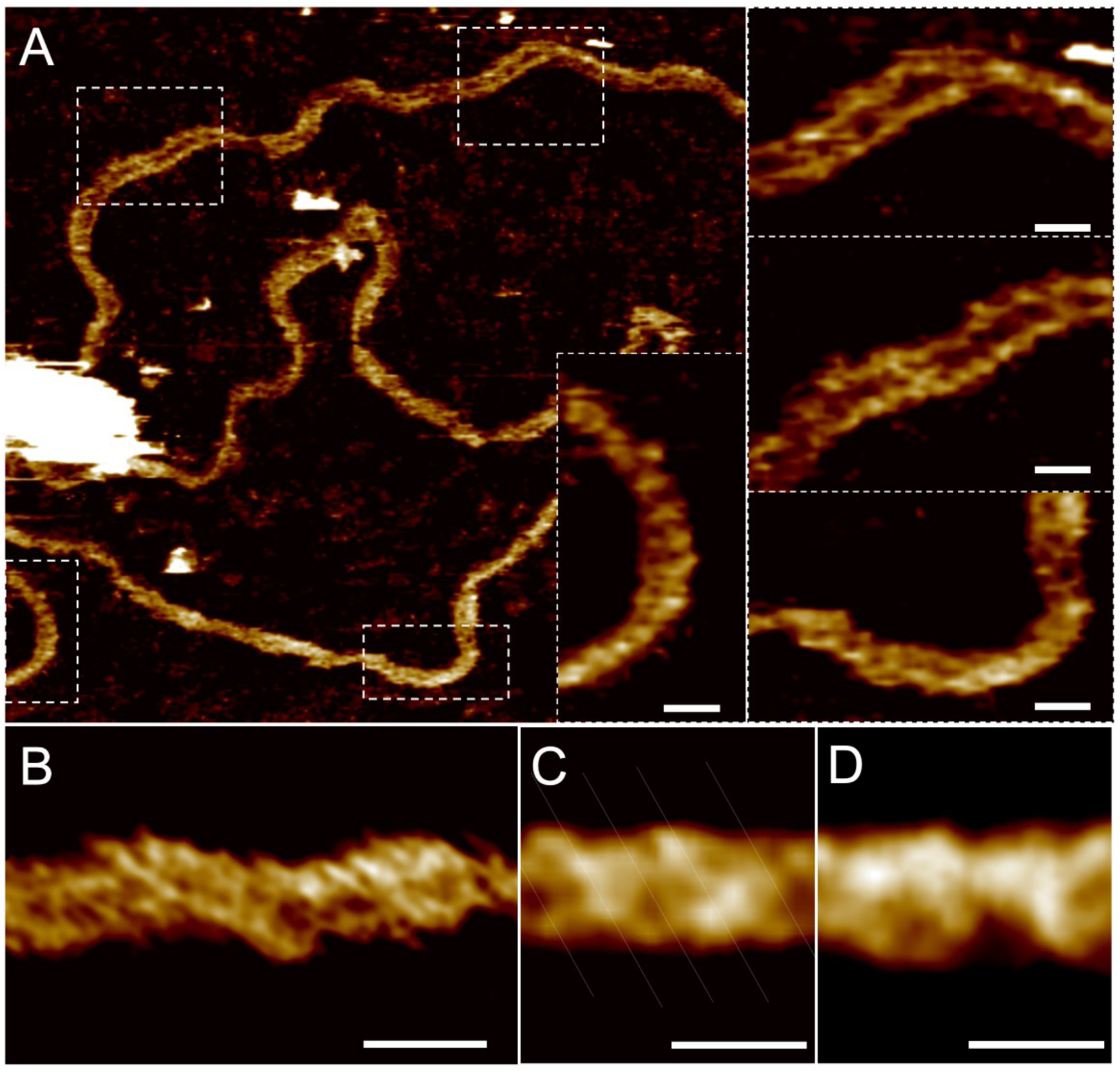
AFM imaging of RNPs on mica in liquid. (**A**) AFM image of RNP with digital zooms displaying helical features. (**B**) High resolution AFM image of RNP. (**C**) Correlation average along the length of a digitally straitened protein using a 20 nm reference section of the RNP in (A) (n = 80). (**D**) Correlation average using the 20 nm reference section with Y-inverted (n = 80). All scale bars: 10 nm.

**Figure S10.**
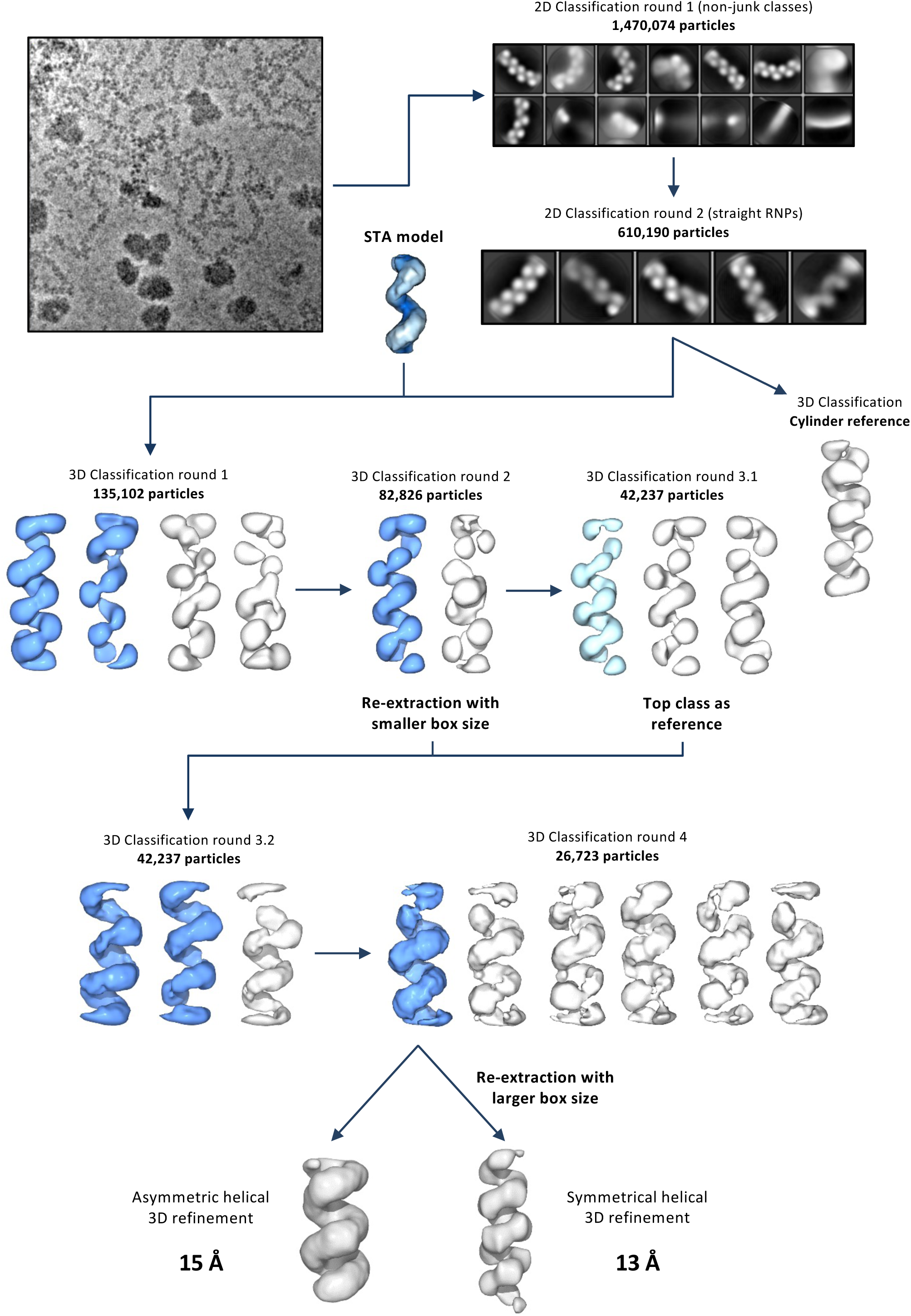
Schematic illustration of the image processing of cryo-EM data. Blue volumes denote the classes taken forwards to further processing.

**Figure S11.**
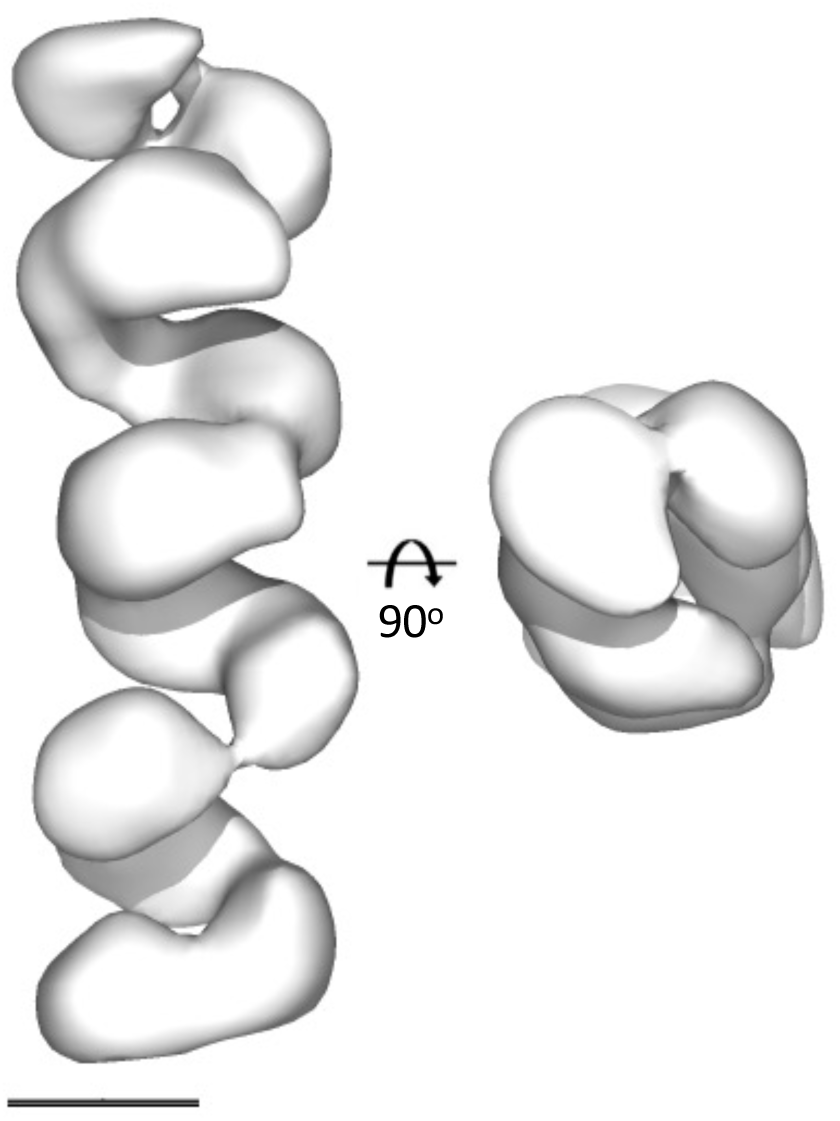
Unbiased processing of BUNV RNP cryo-EM data. Classification with a cylinder reference showed a clear helix in the absence of any symmetry or reference bias.

**Figure S12.**
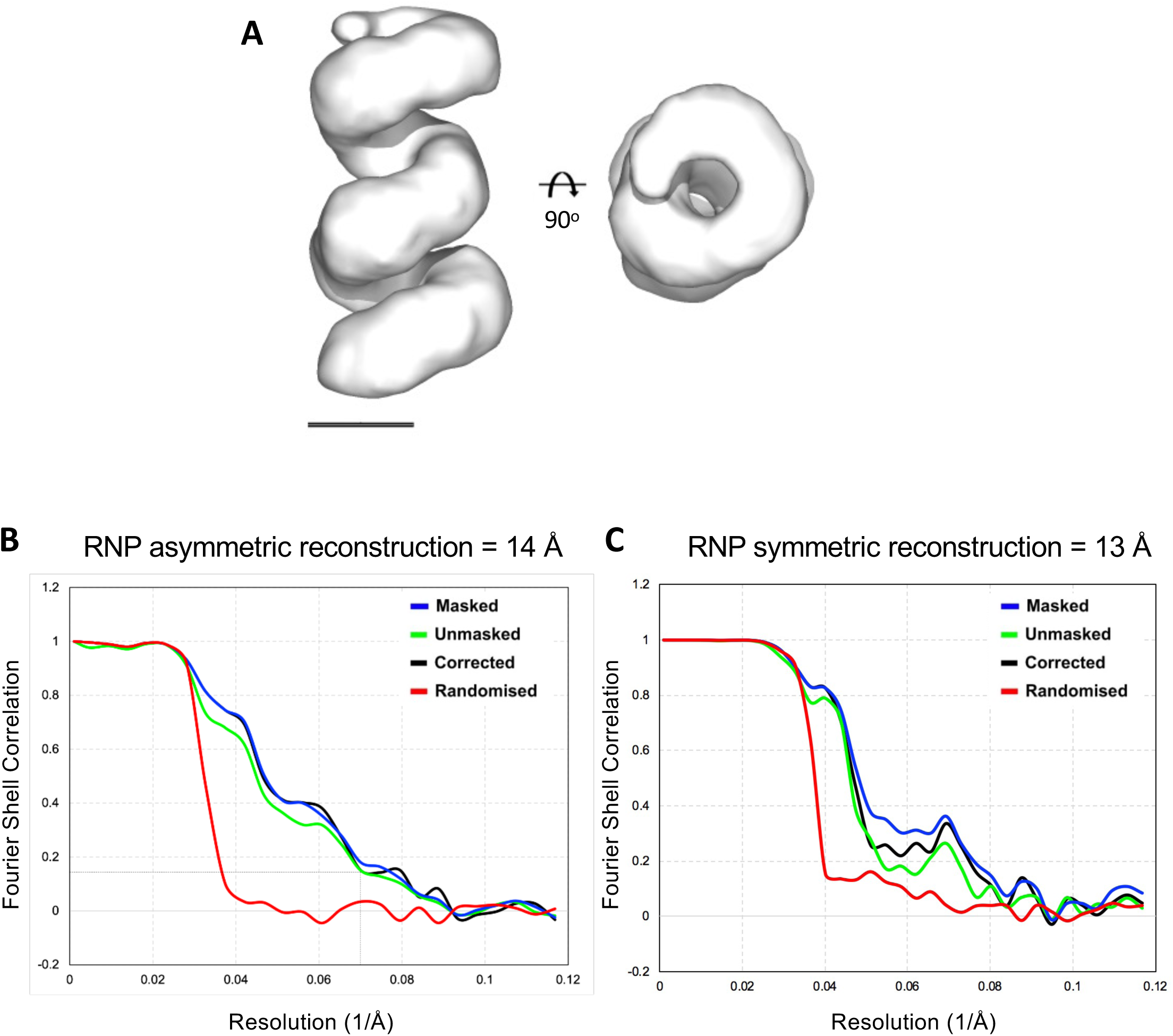
BUNV RNP cryo-EM averages. (**A**) Asymmetric average of a BUNV RNP at 15Å. FSC curves of (**B**) the asymmetric average and (**C**) the symmetric average. FSC cut-off 0.143.

**Figure S13.**
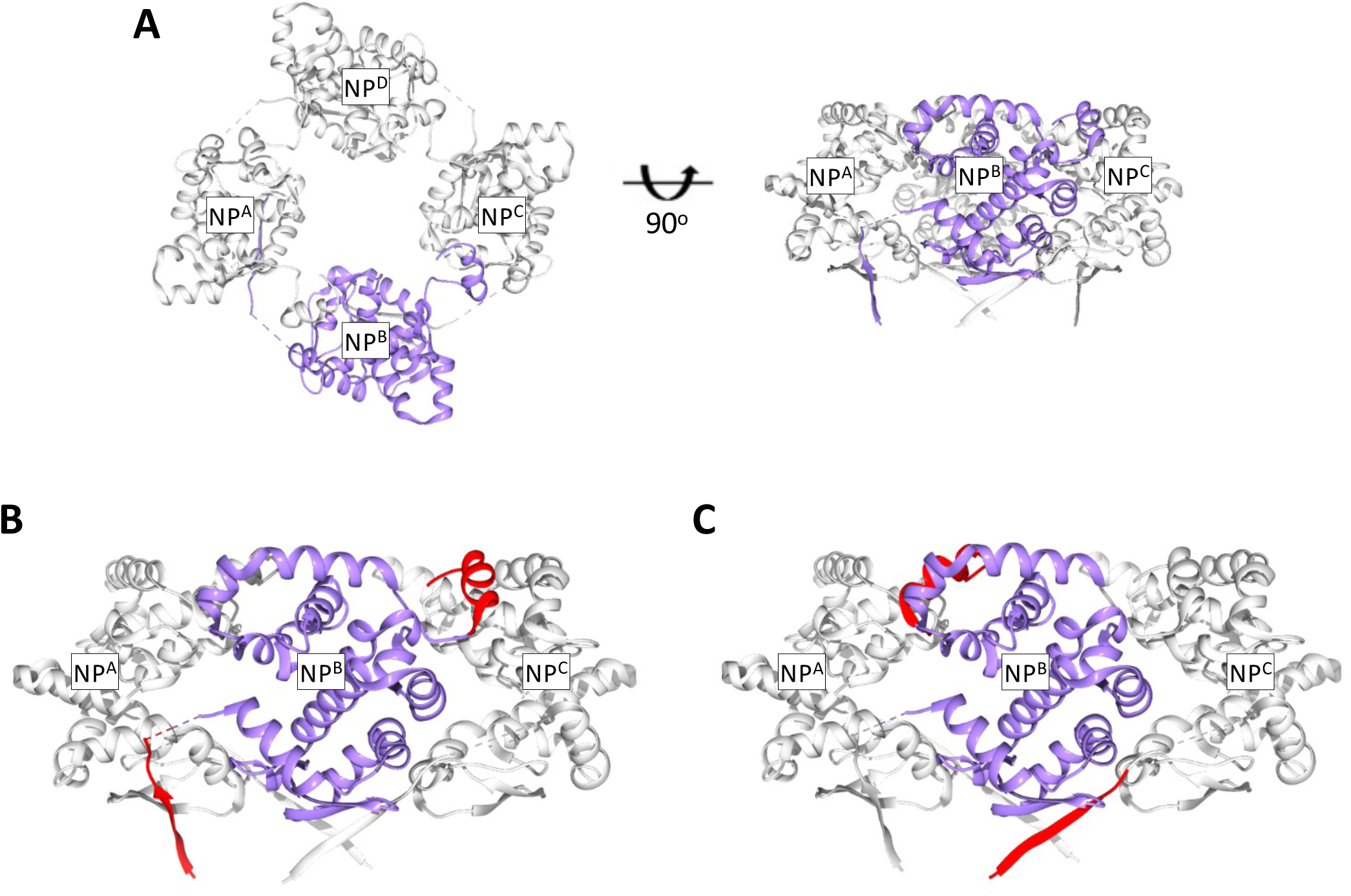
Generation of a ‘split’ BUNV NP for fitting NP to the BUNV RNP model. **(A)** Illustration of the arrangement of NP within an available tetrameric crystal structure. **(B)** A full NP molecule within the crystal structure with the core of NP^B^ in purple and both arms of NP^B^ in red. **(C)** A ‘split’ NP, comprising the core of NP^B^ (purple) and the arms of neighbouring NP^A^ and NP^C^ (red).

**Figure S14.**
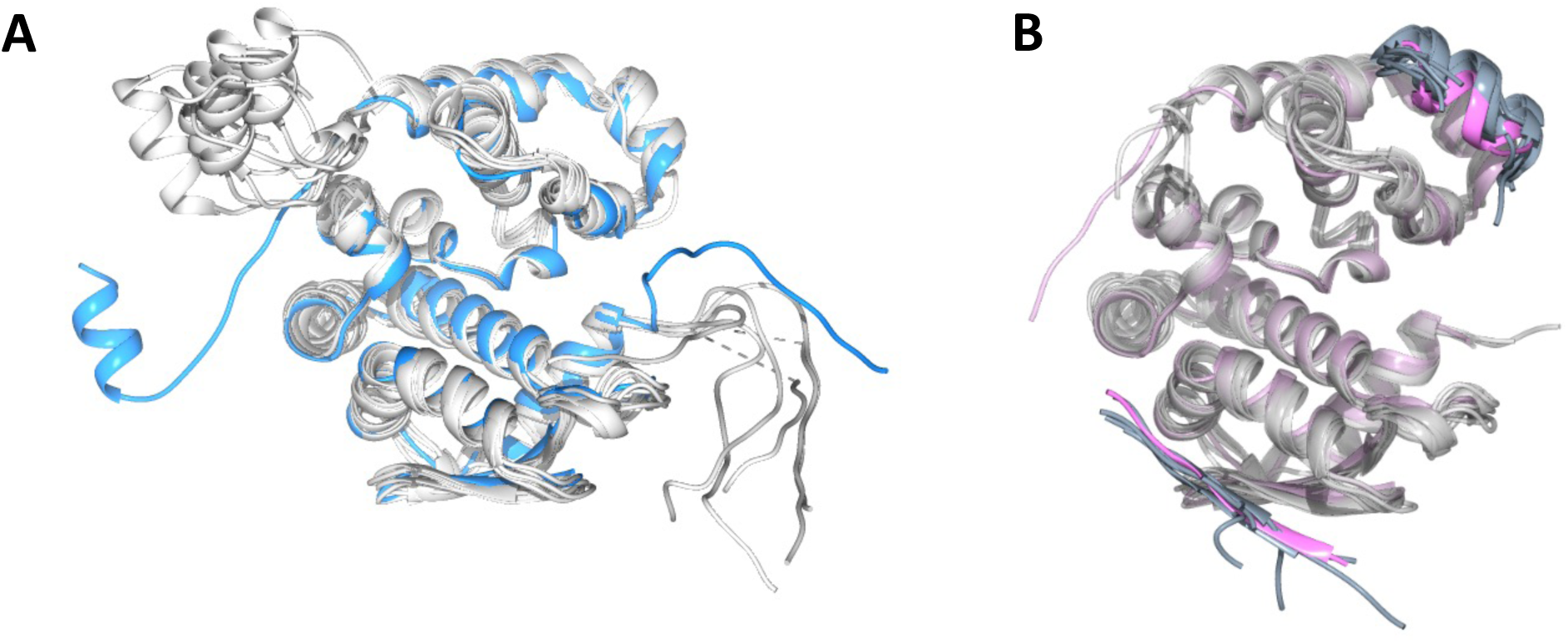
Conservation of the ‘split’ BUNV NP. **(A)** Alignment of available full NP structures from different orthobunyaviruses. Our RNP-derived model in blue. The arms are clearly flexible. **(B)** Alignment of ‘split’ NPs from the same orthobunyaviruses as panel A. Our RNP-derived model in pink. PDB accession codes: 3ZLA, 3ZL9, 4BHH, 4IJS, 4J1J, 4JNG.

**Figure S15.**
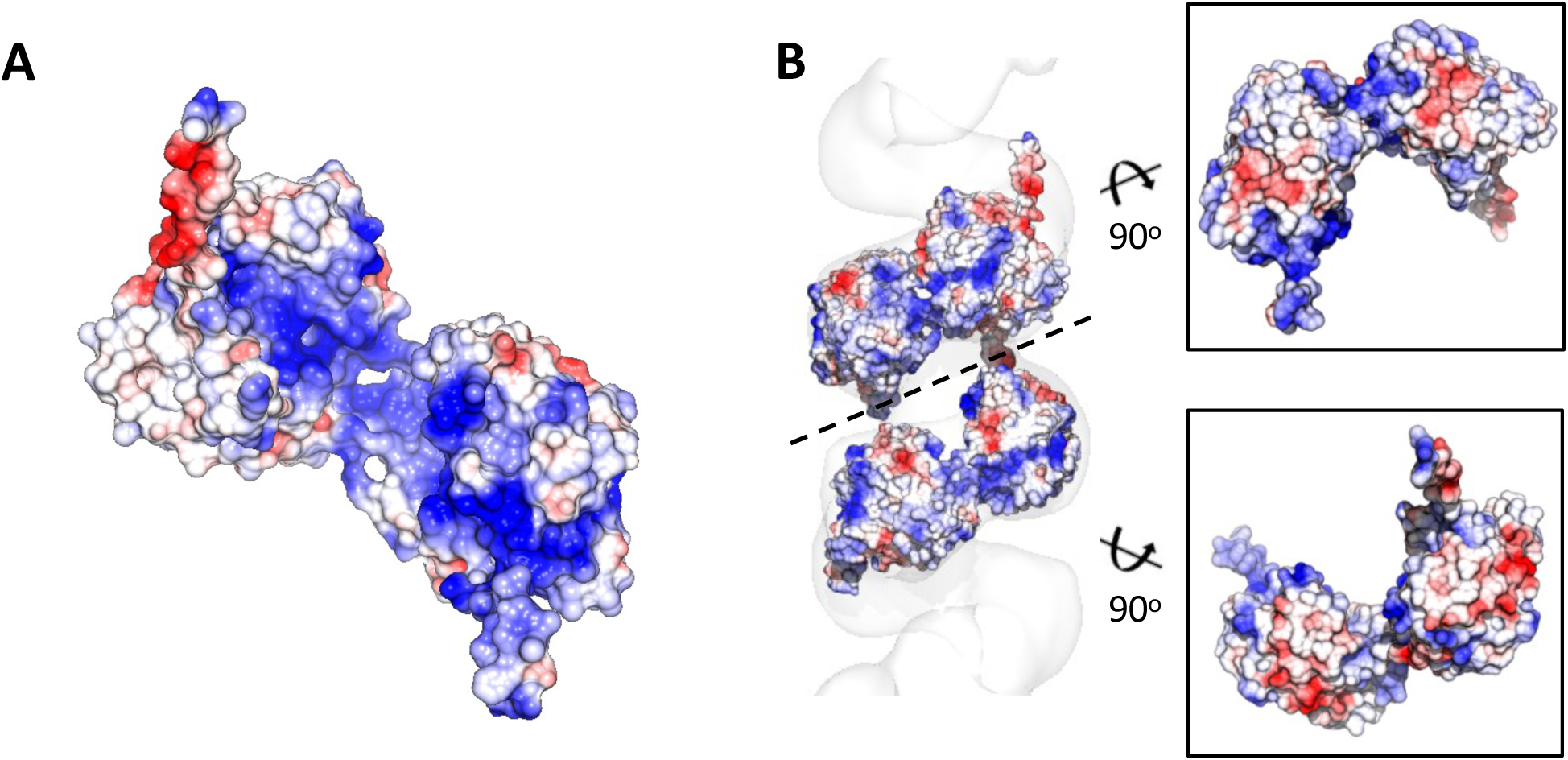
Coulombic surface potential of the helical BUNV NP model. (**A**) The BUNV RNP model has a continuous positively-charged RNA-binding groove pointing towards the inside of the RNP. (**B**) Coulombic surface potential shows electrostatic interactions between lateral neighboring NP monomers are unlikely.

**Figure S16.**
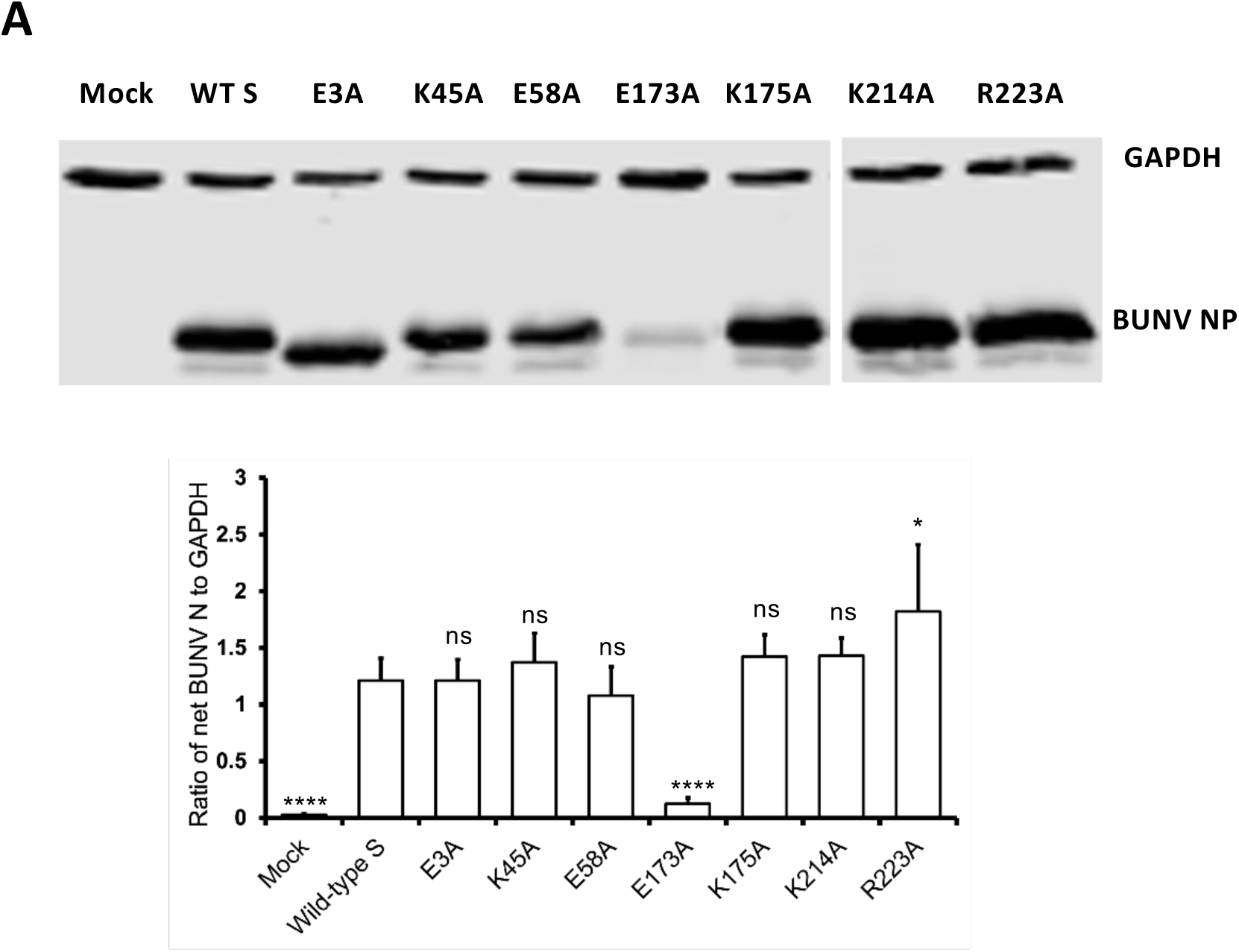
Comparison of expression of mutant compared to wild type NP. ns: p>0.85, *: p ≤ 0.05, **: p ≤0.01, ***: p≤0.001, ****: p≤0.0001.

**Figure S17.**
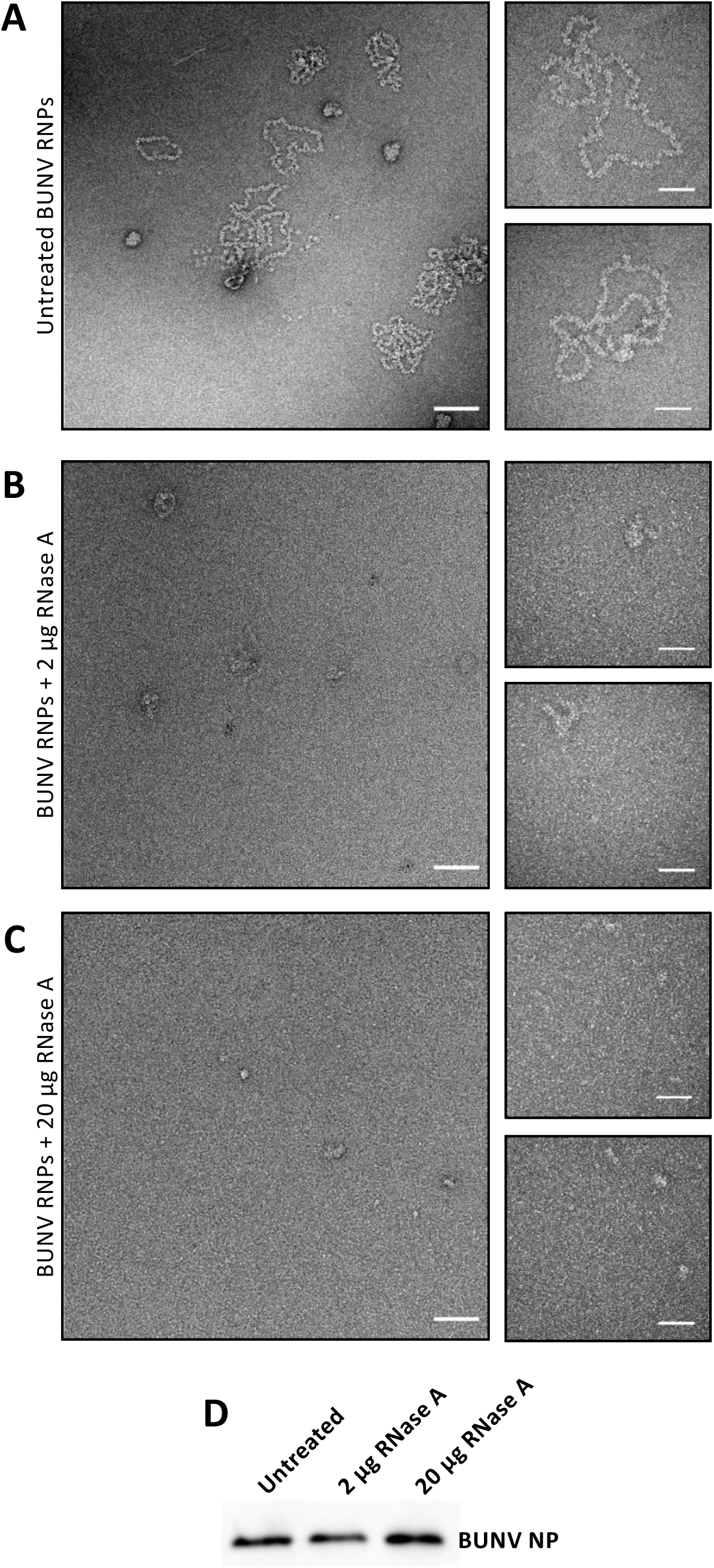
The RNA component of BUNV RNPs maintains their helical arrangement. Purified RNPs were treated with different amounts of RNase A and visualized using negative staining EM. Representative micrographs of (**A**) untreated BUNV RNPs, (**B**) BUNV RNPs treated with 2 μg of RNase A and (**C**) BUNV RNPs treated with 20 μg of RNase A. (**D**) Western blotting against BUNV NP of the different samples imaged by negative stain EM. Scale bars: 100 nm in main and 50 nm in insets.

**Figure S18:**
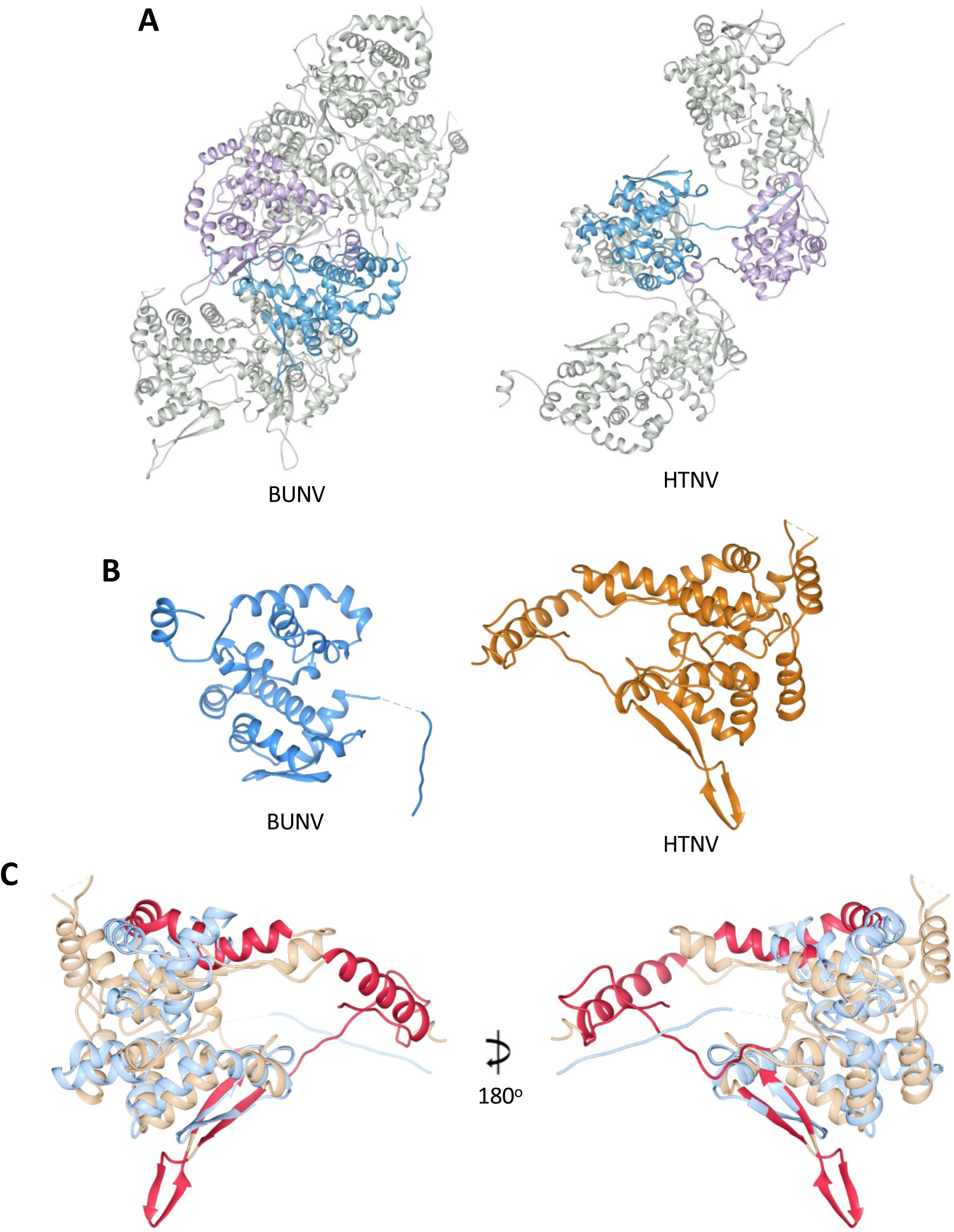
Comparison of BUNV and HTNV RNPs. (**A**) Side by side comparison of helical HTNV and BUNV RNPs. HTNV PDB accession code: 6I2N. (**B**) A side by side comparison of BUNV and HTNV NP. (**C**) Alignment of BUNV (blue) and HTNV (orange) NPs, with HTNV NP homotypic binding residues highlighted in red.

**Table S1:**
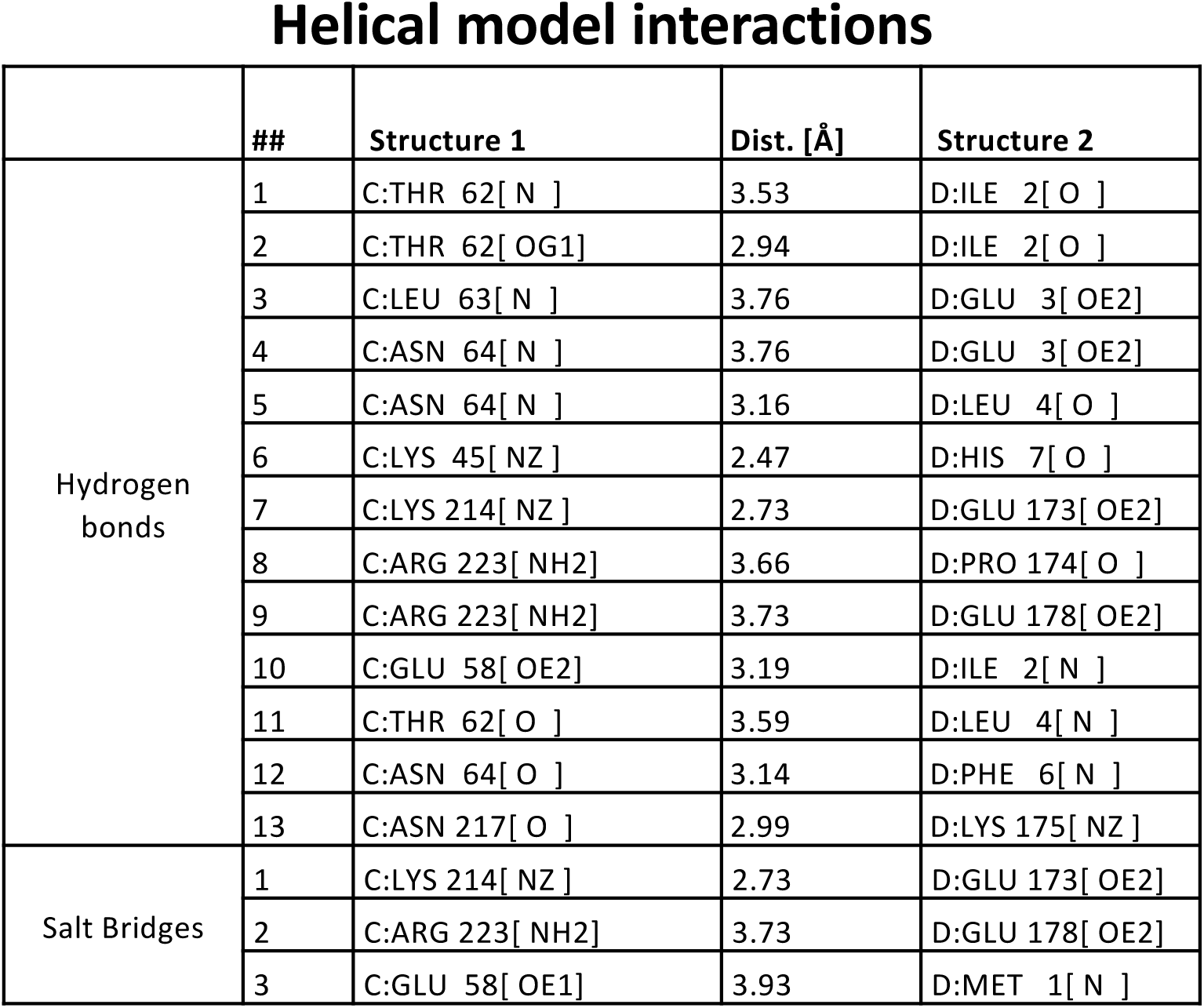
Helical model NP-NP interactions as defined by PDBePISA.

|  | ## | Structure 1 | Dist. [Å] | Structure 2 |
| --- | --- | --- | --- | --- |
| Hydrogen bonds | 1 | C:THR 62[ N ] | 3.53 | D:ILE 2[ O ] |
|  | 2 | C:THR 62[ OG1] | 2.94 | D:ILE 2[ O ] |
|  | 3 | C:LEU 63[ N ] | 3.76 | D:GLU 3[ OE2] |
|  | 4 | C:ASN 64[ N ] | 3.76 | D:GLU 3[ OE2] |
|  | 5 | C:ASN 64[ N ] | 3.16 | D:LEU 4[ O ] |
|  | 6 | C:LYS 45[ NZ ] | 2.47 | D:HIS 7[ O ] |
|  | 7 | C:LYS 214[ NZ ] | 2.73 | D:GLU 173[ OE2] |
|  | 8 | C:ARG 223[ NH2] | 3.66 | D:PRO 174[ O ] |
|  | 9 | C:ARG 223[ NH2] | 3.73 | D:GLU 178[ OE2] |
|  | 10 | C:GLU 58[ OE2] | 3.19 | D:ILE 2[ N ] |
|  | 11 | C:THR 62[ O ] | 3.59 | D:LEU 4[ N ] |
|  | 12 | C:ASN 64[ O ] | 3.14 | D:PHE 6[ N ] |
|  | 13 | C:ASN 217[ O ] | 2.99 | D:LYS 175[ NZ ] |
| Salt Bridges | 1 | C:LYS 214[ NZ ] | 2.73 | D:GLU 173[ OE2] |
|  | 2 | C:ARG 223[ NH2] | 3.73 | D:GLU 178[ OE2] |
|  | 3 | C:GLU 58[ OE1] | 3.93 | D:MET 1[ N ] |

**Table S2.** Tetrameric model (pdb: 3ZLA) NP-NP interactions as defined by PDBePISA.

|  | ## | Structure 1 | Dist. [Å] | Structure 2 |
| --- | --- | --- | --- | --- |
| Hydrogen bonds | 1 | B:LEU 4[ N ] | 3.31 | A:THR 62[ O ] |
|  | 2 | B:PHE 6[ N ] | 2.95 | A:ASN 64[ O ] |
|  | 3 | B:GLN 168[ NE2] | 2.63 | A:PHE 229[ O ] |
|  | 4 | B:ILE 2[ O ] | 3.16 | A:THR 62[ N ] |
|  | 5 | B:ILE 2[ O ] | 3.19 | A:THR 62[ OG1] |
|  | 6 | B:LEU 4[ O ] | 3.05 | A:ASN 64[ N ] |
|  | 7 | B:PHE 6[ O ] | 2.97 | A:GLY 66[ N ] |
|  | 8 | B:ASP 8[ OD2] | 3.39 | A:ARG 40[ NH1] |
|  | 9 | B:GLU 178[ OE2] | 3.50 | A:ASN 217[ N ] |
| Salt Bridges | 1 | B:ASP 8[ OD2] | 3.39 | A:ARG 40[ NH1] |

**Table S3.** BUNV NP helical model NP-NP interactions as defined by PDBePISA. Highlighted in red: mutants tested in the mini-replicon assay.

|  | ## | Structure 1 | Dist. [Å] | Structure 2 |
| --- | --- | --- | --- | --- |
| Hydrogen bonds | 3 | C:LEU 63[ N ] | 3.76 | D:GLU 3[ OE2] |
|  | 4 | C:ASN 64[ N ] | 3.76 | D:GLU 3[ OE2] |
|  | 6 | C:LYS 45[ NZ ] | 2.47 | D:HIS 7[ O ] |
|  | 7 | C:LYS 214[ NZ ] | 2.73 | D:GLU 173[ OE2] |
|  | 8 | C:ARG 223[ NH2] | 3.66 | D:PRO 174[ O ] |
|  | 9 | C:ARG 223[ NH2] | 3.73 | D:GLU 178[ OE2] |
|  | 10 | C:GLU 58[ OE2] | 3.19 | D:ILE 2[ N ] |
|  | 13 | C:ASN 217[ O ] | 2.99 | D:LYS 175[ NZ ] |
| Salt Bridges | 1 | C:LYS 214[ NZ ] | 2.73 | D:GLU 173[ OE2] |
|  | 2 | C:ARG 223[ NH2] | 3.73 | D:GLU 178[ OE2] |
|  | 3 | C:GLU 58[ OE1] | 3.93 | D:MET 1[ N ] |

